# Calpain cleaves the carboxyl terminus of TRPV1 and modulates receptor tachyphylaxis

**DOI:** 10.1101/2025.04.28.651151

**Authors:** Jinyan Jiang, Xiaochen Wang, Jing Yang, Shuwen Gao, Jixuan Xu, Jiao Liu, Yun Wang, Ping Liang, Ying Zhang

**Affiliations:** Department of Neurobiology, School of Basic Medical Sciences and Neuroscience Research Institute; Key Lab for Neuroscience, Ministry of Education/National Health Commission of China, Peking University, Beijing 100191, China; The First Affiliated Hospital and Institute of Translational Medicine, Zhejiang University School of Medicine, Hangzhou 310003, China; Lishui People’s Hospital, The First Affiliated Hospital of Lishui University, Lishui, Zhejiang 323000, China; Center of Medical and Health Analysis, Peking University Health Science Center, Beijing 100191, China; State Key Laboratory of Natural and Biomimetic Drugs, Peking University, Beijing 100191, China

**Author notes:** These authors contributed equally to this work (Jinyan Jiang, Xiaochen Wang, Jing Yang). Correspondence: **Ying Zhang**, PhD, 38 Xueyuan Road, Beijing 100191, China.; **Ping Liang**, MD, PhD, 268 Kaixuan Road, Hangzhou 310029, China.

**Keywords:** TRPV1, calpain, capsaicin, desensitization, internalization

## Abstract

Transient receptor potential vanilloid-1 (TRPV1) plays a critical role in noxious heat sensation and pain hypersensitivity development in chronic pain. Sustained or repeated exposure to capsaicin, a classic TRPV1 agonist, induces TRPV1 desensitization. This partially accounts for the analgesic effect of capsaicin. However, the regulatory mechanisms of TRPV1 desensitization remain poorly understood. In this study, we found that capsaicin acts on TRPV1 to induce the activation of calpain, a Ca^2+^-sensitive cysteine protease, in a manner unrelated to cellular injury. Calpain cleaves rTRPV1 at the carboxyl terminus of G819/S820. Lack of the distal carboxyl terminus leads to reduced TRPV1 localization on the plasma membrane, potentially due to increased receptor internalization and impaired subunit assembly. This finding was corroborated by whole-cell patch clamp recordings. Additionally, the Δ820-838 mutant of rTRPV1 shows resistance to tachyphylaxis as induced by repetitive capsaicin stimulation. This study reveals a pivotal role for calpain in TRPV1 desensitization where its activation constrains TRPV1 channel function while simultaneously increasing its resistance to tachyphylaxis, thereby acting to maintain TRPV1 activity within an appropriate range.

## Introduction

Chronic pain refers to pain that persists for more than three months after the initial injury or disease. It represents the most common cause of years lost to disability^1^. The global prevalence of chronic pain is approximately 30%, suggesting an increased need for safer and more effective analgesic agents. Among numerous receptors and ion channels involved in pain modulation, Transient receptor potential vanilloid-1 (TRPV1) has long been of interest due to its crucial role in pain transmission and modulation. It is closely associated not only with nociceptive heat sensation but also with pain hypersensitivity and the development of peripheral sensitization in chronic pain^2, 3, 4^. These factors highlight TRPV1 as a critical target for analgesic drug development.

TRPV1 is a polymodal nociceptive detector highly expressed in primary sensory neurons. It can be activated by noxious heat (>43°C), protons, endocannabinoids, or other lipid metabolites. The functional TRPV1 channel exists as a tetramer, with its central pore formed by S5-S6 and flanked by S1-S4^5, 6^. Each subunit consists of six transmembrane helices (S1-S6), a pore-forming domain, and intracellular amino (N) and carboxyl (C) termini (hereafter labelled Nt and Ct respectively). Activation of TRPV1 allows the influx of cations, primarily Ca^2+^ and Na^+^. Consequently, the functional sensitization of TRPV1 plays a key role in the enhanced excitability of peripheral sensory neurons. However, development of TRPV1 antagonists for use as analgesic drugs has been hampered by their severe side effect of increased body core temperature. A recent study revealed the structural basis for this side effect and suggested that TRPV1 antagonists targeting regions away from the S4-S5 linker and TRP region may not affect body temperature^7^.

Paradoxically, capsaicin, a classic TRPV1 agonist, has long been used as an analgesic. High concentration capsaicin (8%) patches, for example, have shown persistent analgesic effects on patients with postherpetic neuralgia^8^. This is considered related to the toxic effect of TRPV1 overactivation and subsequent ablation of axonal terminals. Recently, positive allosteric modulators (PAMs), which do not directly activate TRPV1 but enhance the efficacy of TRPV1 agonists such as s-RhTx, have been developed to achieve long-lasting analgesia when coadministered with capsaicin^9^. Additionally, low concentration capsaicin (0.1-0.25%), when used repeatedly, has been shown to reduce the Visual Analogue Scale score of patients with neuropathic pain^10^. This analgesic effect of capsaicin has been associated with TRPV1 desensitization and inactivation of voltage-gated sodium channels or voltage-gated calcium channels. However, due to the initial burning sensation, concurrent anesthesia is usually needed for the application of capsaicin to relieve pain.

Further elucidation of the molecular regulatory mechanisms of TRPV1 could greatly facilitate the development of novel and improved analgesic drugs targeting this channel. To this end we noticed that TRPV1 current exhibits prominent Ca^2+^-dependent desensitization, as characterized by acute desensitization, a rapid reduction of channel activity during a single activator stimulation, and tachyphylaxis, resulting in successive reduced responsiveness to repeated agonist stimulation. From a mechanistic perspective, tachyphylaxis can be partially explained by a failure of recovery from acute desensitization^11, 12^. Multiple mechanisms, including calcineurin dependent dephosphorylation^13^, binding of calmodulin (CaM) to the ankyrin-like repeat domain^14, 15, 16, 17^, and calcium-induced activation of phospholipase C followed by phosphatidylinositol 4,5-bisphosphate (PIP_2_) degradation^18, 19, 20^, have been noted to contribute to such TRPV1 desensitization. In addition, recruitment of the phosphodiersterase PDE4D5 to the plasma membrane by β-arrestin-2 is also required for TRPV1 desensitization^21^. One further study has indicated the magnitude of TRPV1 tachyphylaxis to be inversely proportional to receptor recycling following agonist-induced receptor internalization^22^. In terms of the critical domains of TRPV1 involved in its desensitization, both intracellular N and C termini are required, not only because of the presence of phosphorylational sites, CaM binding, or PIP_2_ interaction sites, but also because of the direct Nt-Ct interaction^23, 24, 25^. As TRPV1 transitions from the activation state to the desensitization state, an interaction between the Nt-Ct occurs, leading to constriction of the channel’s outer pore and subsequent reduction in channel permeability.

Based on previous studies, we postulated that TRPV1 tachyphylaxis might be regulated at the protein level in a Ca^2+^-sensitive manner. Therefore, we investigated the potential role of calpain, a Ca^2+^-activated cysteine protease, in the functional regulation of TRPV1. Our results showed that calpain can be activated by a regular dose of capsaicin. Furthermore, calpain cleaves rTRPV1 at G819/S820, located in the distal Ct. This cleavage not only promoted TRPV1 internalization but also impaired subunit assembly. Notably, the rTRPV1 Δ820-838 mutant (hereafter simply TRPV1 Δ820) showed resistance to tachyphylaxis, although the TRPV1 channel current density is diminished. Altogether, this work reveals calpain, a novel regulator of TRPV1 tachyphylaxis which constrains channel function and concurrently enhances TRPV1 channel resistance to tachyphylaxis.

## Results

### Inhibition of calpain activity reverses the activity-dependent reduction of TRPV1 protein

To determine the effect of receptor activity on TRPV1 protein, we examined the expression level of TRPV1 protein after capsaicin stimulation. For this, N-terminal GST-tag or C-terminal GFP-tag TRPV1 was transiently transfected into human embryonic kidney (HEK) 293 cells. After 24 hours, cells were stimulated with capsaicin (1 μM) or solvent (alcohol) for 10 minutes. A significant decrease of TRPV1 protein was observed in the capsaicin-stimulated cells **(Fig. 1a, b),** whilst no change in the level of cleaved caspase-3 was observed between vehicle- and capsaicin-stimulated cells **(Fig. 1c)**. This indicated the reduction of TRPV1 protein to not be associated with apoptosis. Additionally, we used Chinese hamster ovary (CHO) cells stably expressing TRPV1 (hereafter referred to as CHO-TRPV1 cells) to detect the effect of capsaicin treatment on TRPV1 protein. Downregulation of TRPV1 protein expression was observed 30 minutes after capsaicin (1 μM) stimulation **(Fig. 1d)**. Treatment for shorter time (5 or 10 minutes), failed to induce any significant change in the level of TRPV1 protein **(Supplementary Fig. 1a)**. We suspected that the temporal discrepancy for the agonist-induced diminishment of TRPV1 protein might be attributed to the differential expression levels of TRPV1 in transiently or stably expressing cells. Therefore, CHO-TRPV1 cells were used in the following studies. Moreover, in order to achieve a stable and pronounced decrease of TRPV1 protein, the duration for capsaicin treatment was elongated to 1 hour.

**Fig. 1.**
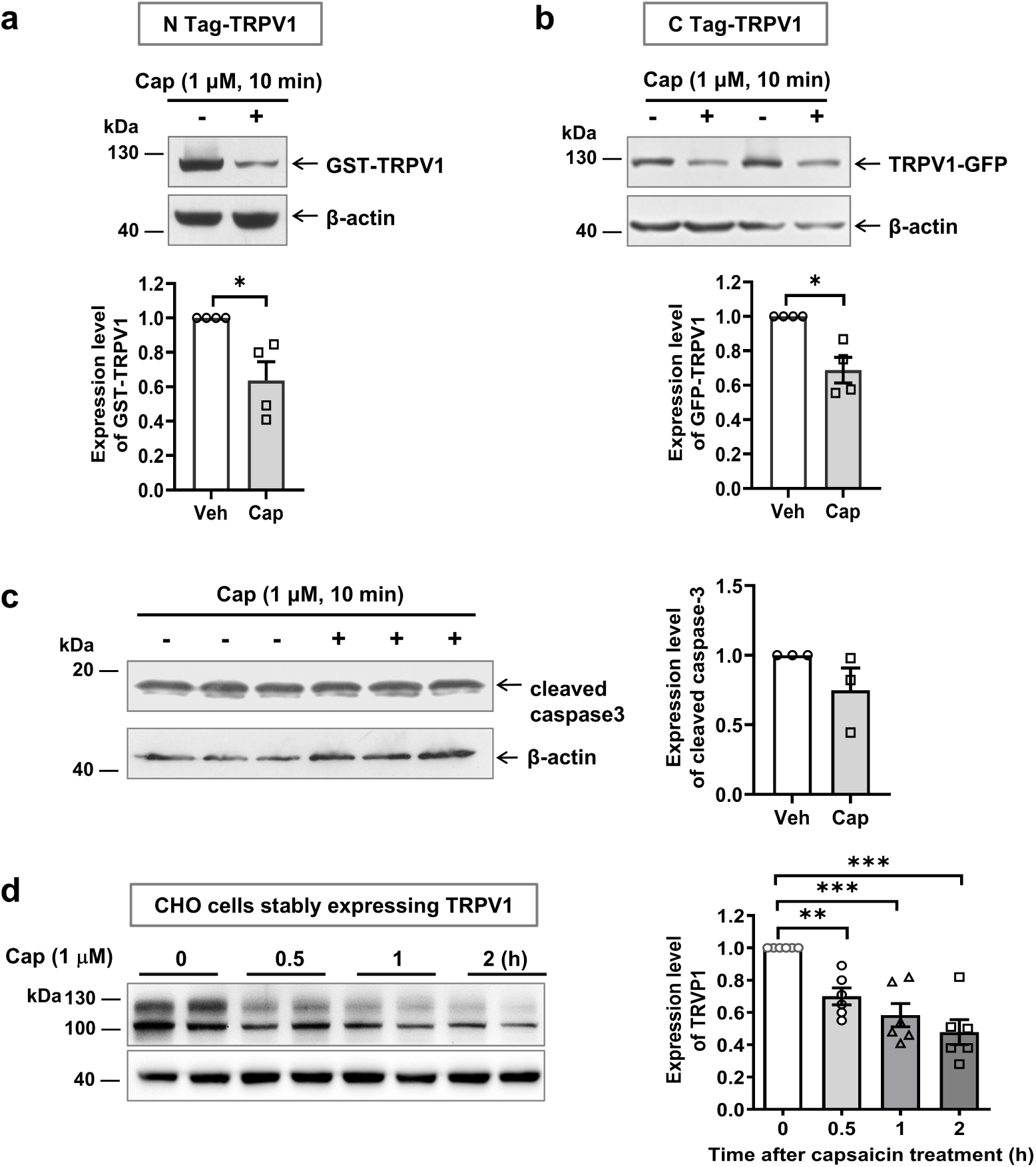
Capsaicin stimulation reduces TRPV1 protein. **a, b.** Treatment with capsaicin (1 μM) for 10 minutes reduced the expression of TRPV1 protein in HEK 293 cells transiently transfected with GST-TRPV (**a**) or TRPV1-GFP (**b**) plasmid. Alcohol, as a solvent, was used as a control. Statistical analysis was performed using unpaired *t*-test with Welch’s correction, n= 4, *\*p*< 0.05. **c.** No change in the expression of cleaved caspase-3 was detected after capsaicin treatment. **d.** Treatment with capsaicin (1 μM) for 30 minutes, 1 hour, or 2 hours in CHO-TRPV1 cells resulted in a diminishment of TRPV1 protein. Statistical analysis was performed using One-way ANOVA followed by the Dunnett’s multiple comparisons test, n= 6, ^\**\**^*p*< 0.01 and ^\*\**\**^*p*< 0.001.

Given the Ca^2+^-permeability of TRPV1 channels and the role of calpain in protein degradation, we examined the effect of calpain inhibitors in reversing the activity-dependent reduction of TRPV1 protein. As expected, pretreatment with distinct calpain inhibitors for 30 minutes, including calpeptin (20 μM), ALLN (20 μM) or MDL28170 (20 μM), significantly reversed or showed a tendency to reverse the capsaicin-induced diminishment of TRPV1 protein **(Fig. 2a, b)**. The effect of the ubiquitin pathway inhibitor MG132 (25 μM) and the caspase inhibitor Z-VAD-FMK (10 μM) was also examined **(Supplementary Fig. 2a**, b). Although no statistical significance was achieved, we could not completely rule out the involvement of ubiquitin or caspase pathways in the diminishment of TRPV1 protein. Notably, transfection of CHO-TRPV1 cells with the endogenous calpain inhibitory protein calpastatin (CAST) plasmid also significantly reversed the activity-dependent diminishment of TRPV1 protein **(Fig. 2c, d)**. Taken together, these data support the involvement of calpain in activity-dependent reduction of TRPV1 protein.

**Fig. 2.**
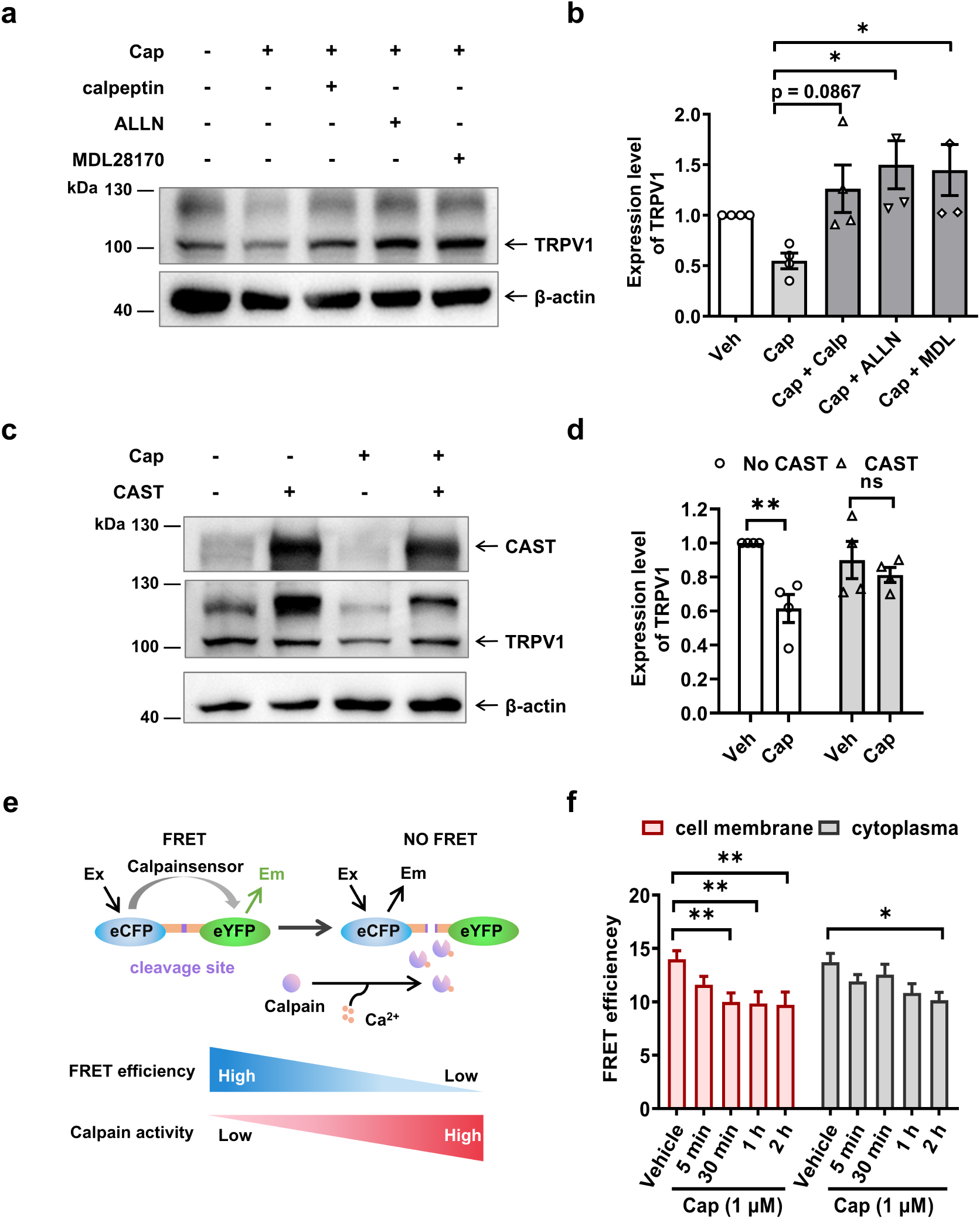
Calpain mediates the activity-dependent decrease of TRPV1 protein. **a, b.** Pretreatment with calpain inhibitor for 30 minutes reversed the decrease in TRPV1 protein in CHO-TRPV1 treated with capsaicin (1 μM, 1 hour). Statistical analysis was performed using One-way ANOVA, n= 4, *\*p*< 0.05. **c, d.** Transfection of CHO-TRPV1 cells with a CAST plasmid reversed the capsaicin-induced reduction of TRPV1 protein. Statistical analysis was performed using two-way ANOVA followed by the Sidak’s multiple comparisons test, n= 4, ns, *p*> 0.05; ^\**\**^*p*< 0.01. **e.** Schematic diagram showing the mechanism of FRET-based calpain activity assay using a calpain sensor. **f.** In CHO-TRPV1 cells treated with 1 μM capsaicin for 30 minutes, a decrease in FRET efficiency was observed near the cell membrane. However, no significant decrease in cytoplasmic FRET efficiency was detected until 2 hours after capsaicin treatment. Statistical analysis was performed using Two-way ANOVA followed by the Sidak’s multiple comparisons test, n= 6, \**p*< 0.05 and ^\**\**^*p* < 0.01. CAST, calpastatin.

Next, we directly measured the activity of calpain following capsaicin treatment. Measurement using a calpain activity assay kit showed that capsaicin (1 μM) treatment for 30 minutes significantly increased the activity of calpain **(Supplementary Fig. 2e)**. To further determine the activity of calpain at the subcellular level, we used fluorescence resonance energy transfer (FRET) acceptor photobleaching to distinguish the activity of calpain near the plasma membrane or in the cytoplasm. A decrease in FRET efficiency denoted an increase of calpain activity **(Fig. 2e)**. We observed that calpain activity near the plasma membrane showed a tendency to decline 5 minutes after capsaicin treatment, with statistical significance achieved at 30 minutes **(Fig. 2f and Supplementary Fig. 2c)**. The calpain activity in the cytoplasm was also significantly decreased 2 hours post capsaicin treatment. However, no significant change in cell morphology was observed 2 hours post capsaicin treatment **(Supplementary Fig. 2d)**, suggesting the calpain activation in the present study to lack the widespread cytotoxic effects that accompany the use of capsaicin.

### Calpain cleaves TRPV1 Ct at G819/S820 site

To examine the potential action of calpain on the intracellular fragments of TRPV1, we induced the expression of TRPV1 Nt and TRPV1 Ct *in vitro*. Calpain-1 (μ-calpain, 2.5 U/ml) was directly added to the protein solution and incubated for 30 minutes at 37°C. GST was used as a control. Both the GST-TRPV1 Nt and GST-TRPV1 Ct showed a reduction in protein levels, but not GST *per se* **(Fig. 3a)**. Meanwhile, a remarkable cleavage band was observed in the GST-TRPV1-Ct after incubation with calpain-1. Moreover, the appearance of this cleavage band could be blocked by the calpain inhibitors calpeptin (20 μM) or MDL28170 (20 μM), indicating the specificity of calpain-mediated cleavage **(Fig. 3b)**. This cleavage band of GST-TRPV1 Ct could be detected by an anti-GST antibody **(Fig. 3c, left panel)** but not by an antibody against the TRPV1 Ct (residues 824-838) **(Fig. 3c, right panel)**, indicating that the carboxyl-terminal end of the TRPV1 Ct was deleted by calpain. Combined with the shift in molecular weight of the cleavage fragment, we postulate that calpain might cleave the distal carboxyl end of the TRPV1 Ct.

**Fig. 3.**
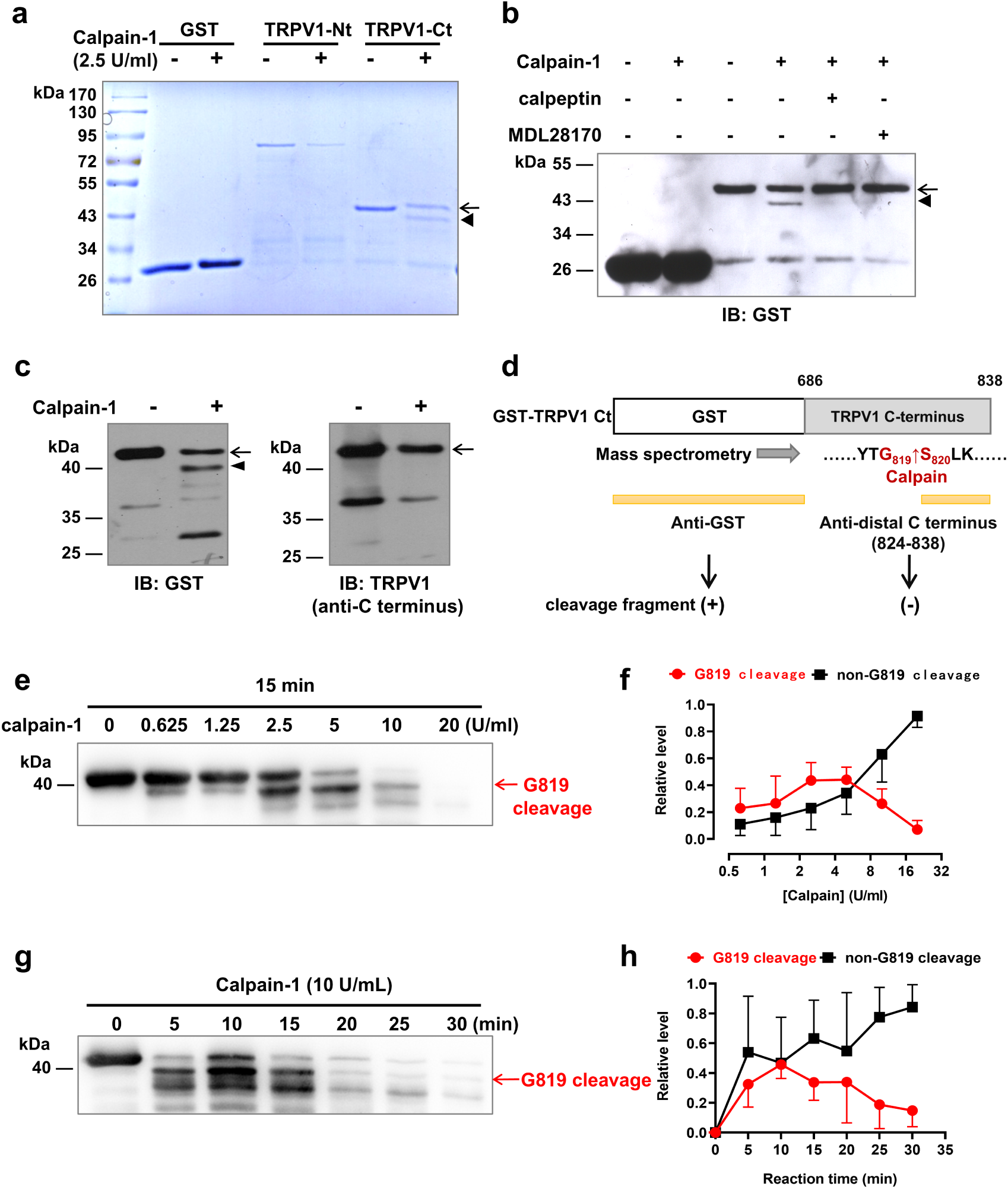
*In vitro* cleavage of TRPV1 Ct by calpain-1. **a.** *In vitro* cleavage of GST, GST-TRPV1 Nt and GST-TRPV1 Ct by calpain-1 (2.5 U/ml). Coomassie brilliant blue staining was used to display the protein bands. The arrow denotes the full-length band of the GST-TRPV1 Ct and the arrowhead denotes the cleavage fragment. **b.** Inhibition of the calpain-mediated cleavage fragment by the calpain inhibitors calpeptin (10 μM) and MDL28170 (10 μM). **c.** Comparison of the results of Western blotting using anti-GST antibody (left) and anti-TRPV1 Ct antibody (right). **d.** Schematic diagram showing the antigenic epitope of anti-GST and anti-TRPV1 Ct (residues 828-838) antibodies, respectively. **e-h.** Time-(**e, f**) and concentration-(**g, h**) dependent cleavage of TRPV1 Ct by calpain. Arrows denote the cleavage fragment. The amount of G819 cleavage was quantified by “G819 cleavage / GST-TRPV1 Ct_0_”. Ct_0_ means that time or the enzyme concentration equals zero. The amount of unspecific cleavage was quantified by “(GST-TRPV1 Ct_0_ – GST-TRPV1 Ct_n_ – G819 cleavage) / GST-TRPV1 Ct_0_”. Ct_n_ refers to the indicated reaction time or enzyme concentration.

To precisely determine the cleavage sites of calpain on TRPV1 Ct, we excised the differential protein bands in the SDS-PAGE gel of TRPV1 Ct subjected to calpain treatment for mass spectrometry analysis **(Supplementary Fig. 3a, GST-TRPV1 Ct and Cleavage-1, -2, -3, and -4)**. G819-terminated peptide was detected in Cleavage-1 and -2, but not in GST-TRPV1 Ct or Cleavage-3 or -4 **(Supplementary Fig. 3c-g)**. This finding indicated that the break point at G819 was caused by the action of calpain, but not by sample digestion treatment in the mass spectrometry assay. The presence of G819-terminated peptide in Cleavage-2 was considered potentially to be due to sequential cleavage of TRPV1 Ct following cleavage of the G819/S820 site. In addition, we used GPS-CCD (Calpain Cleavage Detector), a computational program for the prediction of calpain cleavage sites, to predict the possible cleavage site in TRPV1 Ct **(Supplementary Fig. 3b)**^26^. The G819/S820 site was listed as the cleavage site closest to the C-terminal end. No other specific sites were detected in Cleavage-3 and -4, implying that these bands might be attributed to the non-specific cleavage activity of calpain. Additionally, such GS sites are noted as highly conserved in TRPV1 from the mammals listed in **Supplementary Fig. 4**. Combining this with the above data, we focused on the action of calpain on the G819/S820 site of the TRPV1 Ct in the following investigations.

To investigate the optimal conditions for calpain-1-mediated cleavage of TRPV1 Ct, we studied the influence of incubation time and enzyme concentration on the reaction. When incubated for 15 minutes, the optimal concentration of calpain for the production of the G819 cleavage fragment was approximately 2.5 U/ml **(Fig. 3e, f)**. When the calpain concentration was set at 10 U/ml, the optimal incubation time was 10 minutes **(Fig. 3g, h)**. When calpain concentration or reaction time continued to increase, the amount of G819/S820 cleavage fragment gradually reduced until such fragment was completely degraded. This could be attributed to the nonspecific cleavage action of calpain.

### Full-length TRPV1 is cleaved at G819/S820 site by calpain

To validate whether calpain-mediated cleavage occurs in full-length TRPV1, we inserted a Flag tag (DYKDDDDK) at the N-terminus of the TRPV1-GFP plasmid and constructed a Flag-TRPV1-GFP plasmid **(Fig. 4a)**. Transfection of the Flag-TRPV1-GFP plasmid in HEK 293 cells verified the expression of Flag-TRPV1-GFP fusion protein, as detected by anti-Flag or anti-GFP antibody **(Fig. 4b)**. After protein enrichment by immunoprecipitation (IP) with GFP antibody, the protein concentrate was treated with calpain-1 (10 U/ml). Both coomassie brilliant blue staining of SDS-PAGE gel **(Fig. 4c)** and Western blotting **(Fig. 4d)** resulted in the appearance of new bands after calpain treatment, including both of truncated Flag-TRPV1 and GFP-tagged C-terminal fragment. The molecular weight of these bands was conformed with Flag-TRPV1 Δ820 and 820-838-GFP. Regrettably, the bands of truncated TRPV1 could not be specifically recognized by anti-Flag antibody. Accordingly, we performed concentration-dependent cleavage of TRPV1-GFP by calpain-1. Immunoblotting with anti-TRPV1 Ct antibody showed that the potential bands of TRPV1 Δ820 **(Fig. 4e)** and 820-838-GFP **(Fig. 4f)** exhibited a dose-dependent increment as the concentration of calpain-1 increased. These data support the conclusion that calpain cleaves full-length TRPV1 at G819/S820 site in the distal Ct.

**Fig. 4.**
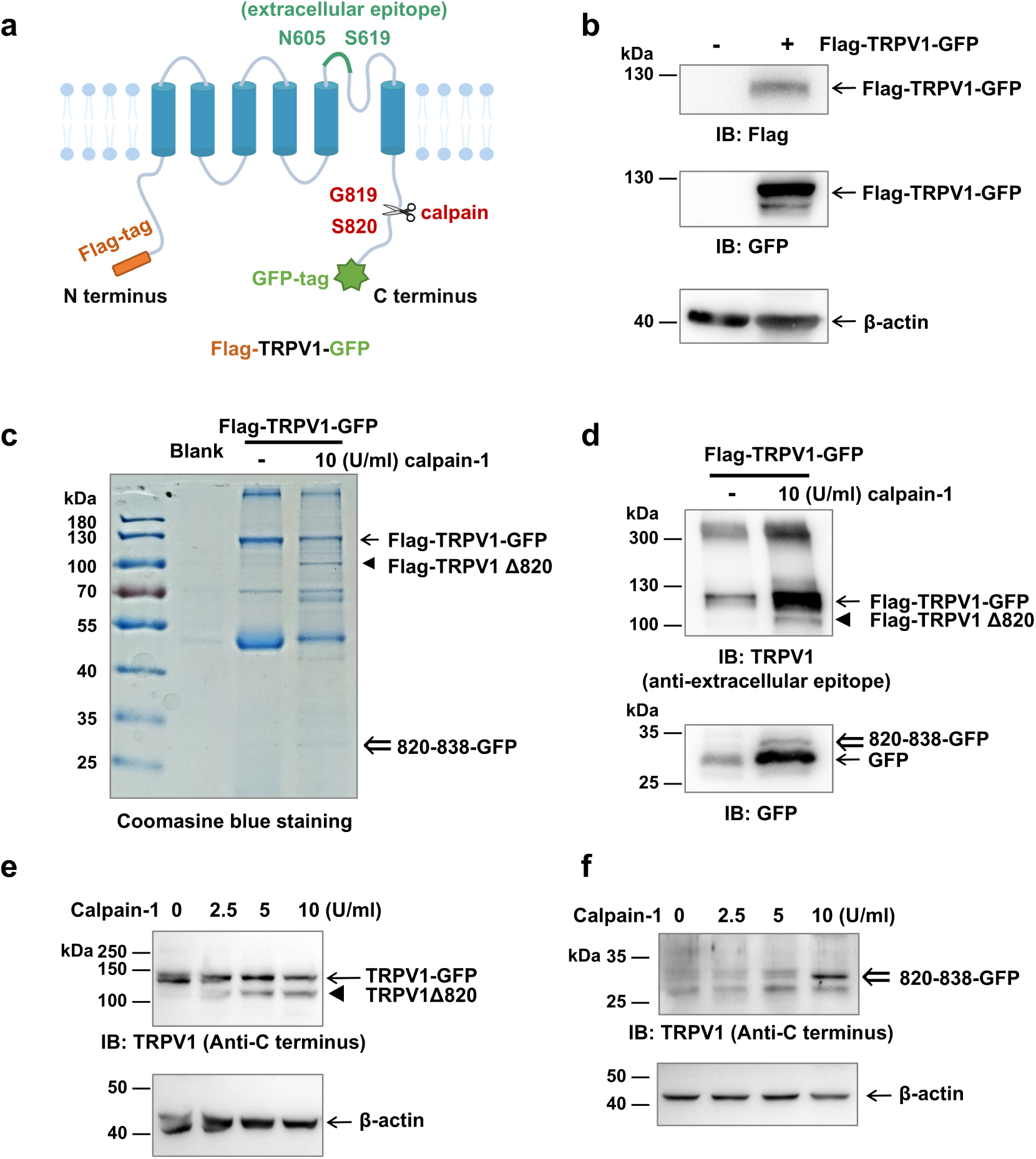
Calpain cleaves full-length TRPV1 potentially at G819/S820 site. **a.** A schematic diagram showing construction of Flag-TRPV1-GFP fusion protein and the potential cleavage sites by calpain. **b.** Expression of Flag-TRPV-GFP plasmid in HEK 293 cells. Protein with its corresponding prediction of molecular weight was detected with anti-Flag and anti-GFP antibodies, respectively. **c.** Gel staining image by coomassie brilliant blue of Flag-TRPV1-GFP (122 kDa) with or without calpain-1 treatment. The potential bands of truncated Flag-TRPV1 Δ820 (93 kDa) and 820-838-GFP (29 kDa) are indicated. **d.** Western blotting of full-length TRPV1 with or without calpain treatment. To present the bands after cleavage more clearly, the amount of protein subject to calpain treatment was 2 times of that without calpain treatment. **e, f.** Concentration-dependent cleavage of TRPV1-GFP by calpain-1. The potential bands of truncated TRPV1 Δ820 (93 kDa, **e**) and 820-838-GFP (**f**) are indicated. “←” denotes full-length TRPV1; “◄” denotes the potential truncated TRPV1 Δ820; “⇐” denotes the cleavage fragment of 820-838-GFP.

To further determine the critical amino acid residues involved in the action of calpain-mediated cleavage, we conducted point mutation studies of GST-TRPV1 Ct. A series of mutants, namely double mutant, triple mutant, quadruple mutant-1, and quadruple mutant-2, were constructed **(Supplementary Fig. 5a)**. However, none of these mutants had a significant impact on the calpain-mediated cleavage **(Supplementary Fig. 5b-d)**. This lack of impact might be caused by the feature of calpain where it recognizes substrates not only by the primary sequence of their amino acids, but also by their overall three-dimensional structure^27^. Further experiments are therefore required to identify the key recognition sites of calpain for the TRPV1 Ct.

### The TRPV1 Δ820 mutant exhibits reduced cell membrane localization

To investigate the potential role of distal Ct of TRPV1 (820-838 amino acid residues) in the regulation of receptor function, we constructed a GFP-tagged truncated mutant of TRPV1, TRPV1 Δ820-GFP **(Supplementary Fig. 5a).** TRPV1-GFP was used as a control. Effective expression of TRPV1 Δ820-GFP in HEK 293 cells was confirmed by Western blotting **(Supplementary Fig. 5b)**. A small shift in protein bands to a lower molecular weight was observed in TRPV1 Δ820-GFP compared with TRPV1-GFP. Furthermore, immunoblotting with anti-TRPV1 Ct antibody examined the expression of TRPV1-GFP, but not TRPV1 Δ820-GFP in which the antibody epitope was deleted. The cellular distribution of TRPV1-GFP and TRPV1 Δ820-GFP was then observed in transfected HEK 293 cells. TRPV1-GFP exhibited significant membrane localization, with green fluorescence forming a circumferential distribution along the plasma membrane **(Fig. 5a, left panel)**. Limited colocalization of green and red fluorescence and the ER Tracker was observed. By contrast, TRPV1 Δ820-GFP did not show obvious membrane localization, instead being diffusely distributed throughout cytoplasm and highly colocalized with ER Tracker. Quantification analysis of the GFP fluorescence intensity spectrum across the cells showed prominent double peaks in the TRPV1-GFP group **(Fig. 5a, right panel)**, unobserved in the GFP or TRPV1 Δ820-GFP groups.

**Fig. 5.**
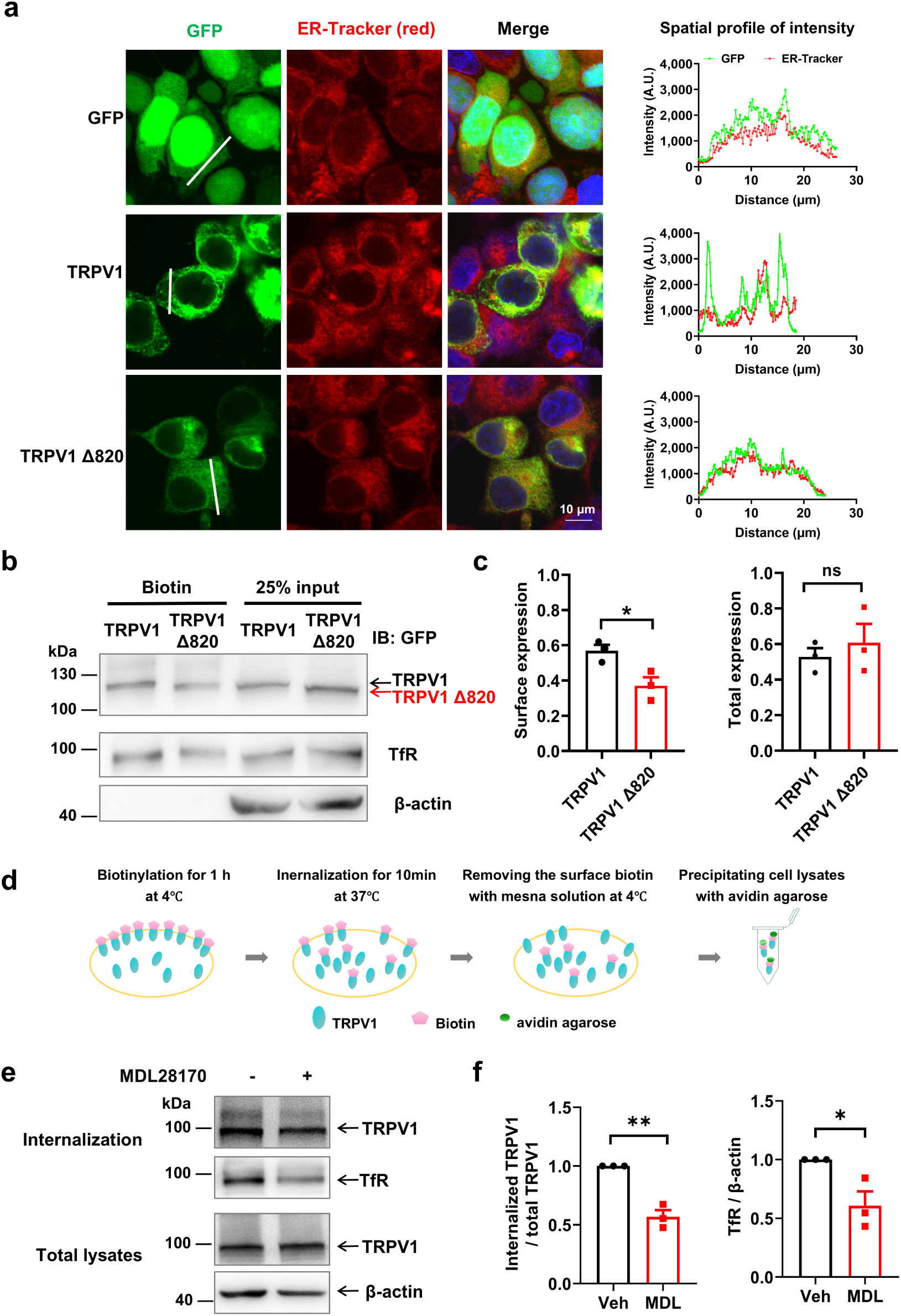
Cleavage of the TRPV1 at G819/S820 site reduces the receptor localization on the cell membrane. **a.** Left, Colabeled images of GFP, TRPV1-GFP or TRPV1 Δ820-GFP and ER Tracker, a marker of endoplasmic reticulum (ER). Right, spatial fluorescence intensity spectrum, with green lines and red lines representing the fluorescence intensity of GFP or ER Tracker, respectively. **b.** Biotinylation assays of the surface level of TRPV1-GFP or TRPV1 Δ820-GFP. Cell surface protein was biotinylated and purified using avidin beads. 25% input was used as a control. Arrows denote TRPV1 or TRPV1 Δ820. **c.** Quantification analysis of the cell surface or total expression levels of TRPV1 or TRPV1 Δ820, with the former normalized to TfR and the latter normalized to β-actin. Statistical analysis was performed using unpaired *t*-tests, n= 3, *\*p*< 0.05. **d** Schematic diagram showing the method to examine receptor internalization. **e.** Effects of calpain inhibition on the constitutive internalization of TRPV1. **f.** Quantification analysis of the internalized receptor. The amount of internalized TRPV1 was quantified as the ratio of internalized TRPV1 to TRPV1 in the total cellular lysates. The amount of internalized TfR was calculated as the ratio of internalized TfR to β-actin. Statistical analysis was performed using unpaired *t*-test, n= 3, *\*p*< 0.05 and ^\**\**^*p*< 0.01.

In addition, a biotin labeling experiment of the cell surface protein was conducted. Results showed that the surface expression of the TRPV1 Δ820 mutant was significantly reduced compared to that of full-length TRPV1 **(Fig. 5b, c)**, despite their expression in cellular lysates did not differ. Transferrin receptor (TfR) was then used as an internal reference for the loading of cell surface protein. Effective separation of the cell surface protein and cytoplasmic protein was demonstrated via the lack of the β-actin bands in the fraction of cell surface protein. These findings suggest that the membrane localization of TRPV1 is compromised upon deletion of the distal Ct, which may then lead to receptor retention in ER.

### Calpain activation promotes TRPV1 internalization and impairs subunit assembly

Combining the above data, we propose that calpain mediates the production of TRPV1 Δ820 and that this truncated form of TRPV1 exhibits reduced cell membrane localization. We therefore assumed that the cell membrane localization of TRPV1 might be regulated by the activity of calpain. As the amounts of receptors on the cell membrane is determined by the relative level of receptor trafficking and receptor internalization, we observed the effects of calpain activity on the constitutive internalization of TRPV1. For this, the biotinylation labeling method conjugated with application of 2-mercaptoethanesulfonic acid (MESNA) was adopted **(Fig. 5d)**. Constitutive internalization of TRPV1 was significantly reduced by the addition of MDL28170 (20 μM) **(Fig. 5e, f)**, suggesting that inhibition of calpain activity had led to a reduction in TRPV1 internalization. Intriguingly, internalization of TfR was also inhibited by MDL28170. From these findings, we could conjecture that calpain-mediated cleavage of TRPV1 may promote receptor internalization.

Previous lines of evidence have shown that functional TRPV1 is localized on the cytomembrane as a tetramer and that the TRPV1 Ct mediates subunit assembly^28^. We therefore determined the effect of Δ820 truncation on the subunit assembly of TRPV1 as a prerequisite for receptor trafficking. To investigate whether removal of the distal Ct of S820 to K838 would affect subunit assembly and cell membrane localization of TRPV1, we conducted an *in vitro* GST-pull down assay and growth associated protein 43 (Gap43) fragment-mediated membrane localization experiment. The GST-pull down assay showed that the level of precipitated His_6_-TRPV1 Ct by GST-TRPV1 Ct Δ820 was significantly lower than that precipitated by GST-TRPV1 Ct **(Fig. 6a, b)**. 25% input was used as a positive control. In addition, we constructed Gap43-TRPV1 Ct, TRPV1 Ct-GFP, and TRPV1 Ct Δ820-GFP plasmids **(Fig. 6c)**. Gap43 has a membrane-targeting segment of 20 amino acid residues at the N-terminus. This was used in the current studies where TRPV1 Ct becomes tethered to the cell membrane when it is fused with Gap43^28^. Western blotting experiments demonstrated that all these plasmids were correctly expressed in HEK 293 cells. As expected, the protein band of TRPV1 Ct Δ820-GFP was slightly lower than that of TRPV1 Ct-GFP. When TRPV1 Ct-GFP or TRPV1 Ct Δ820-GFP was expressed alone, the fluorescence signal remained diffusely distributed throughout the cell and with no obvious fluorescence signal remaining on the cell membrane **(Fig. 6d, left)**. However, when co-transfected with Gap43-TRPV1 Ct, clear fluorescence signals were observed on the cell membrane in the TRPV1 Ct-GFP group, but not in the TRPV1 Ct Δ820-GFP group. Quantification analysis revealed a dual peak signal in the spatial fluorescence intensity spectrum in the TRPV1 Ct-GFP group **(Fig. 6d, right)**, which reflected the direct interaction between Gap43-TRPV1 Ct and TRPV1 Ct-GFP. However, TRPV1 Ct-mediated interaction was compromised by deletion of the distal 19 amino acid residues.

**Fig. 6.**
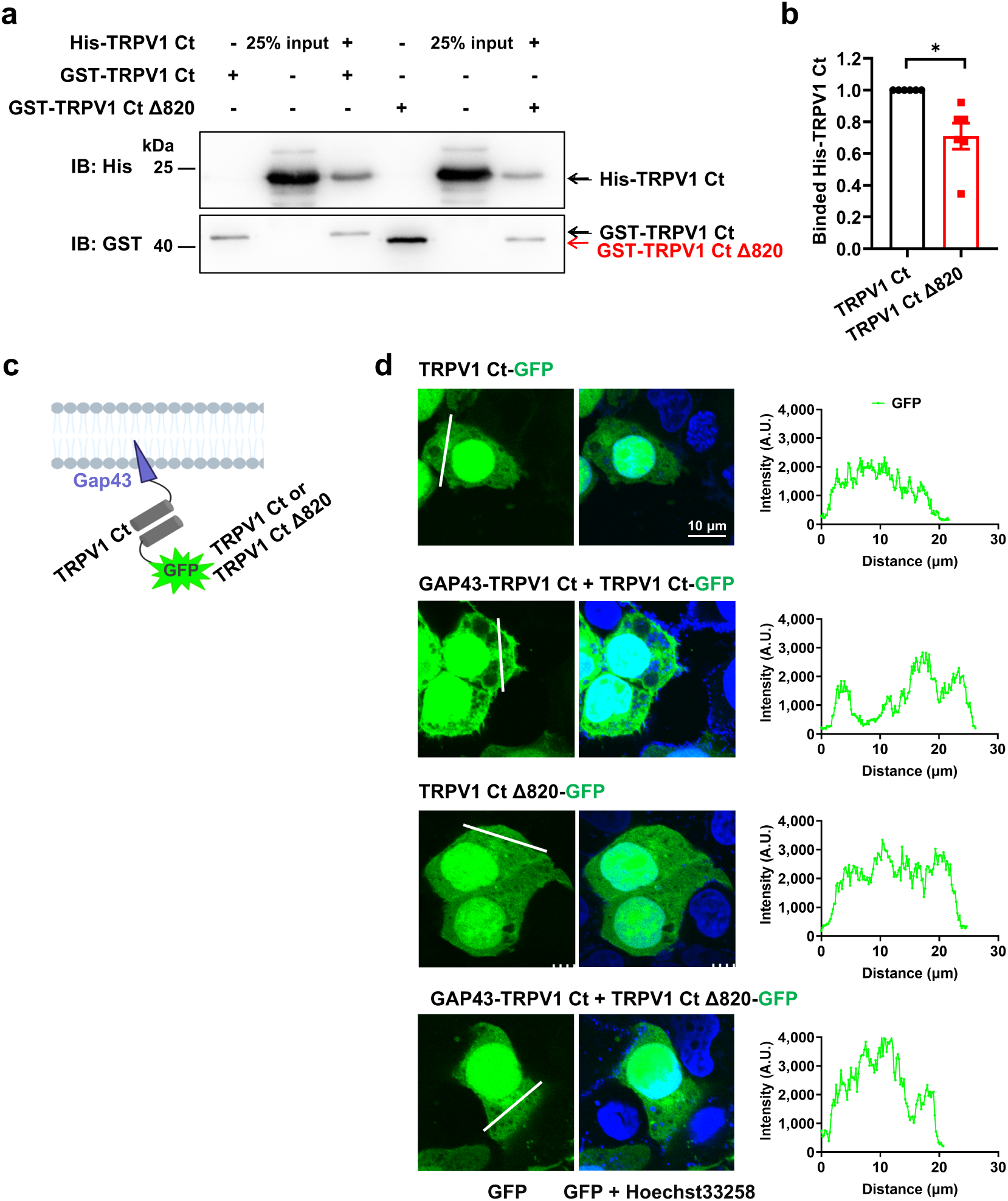
Deleting the distal 19 amino acid residues of TRPV1 Ct impairs subunit assembly. **a.** The *in vitro* GST pull-down experiment showed that substituting GST-TRPV1 Ct with the mutant GST-TRPV1 Ct Δ820 significantly reduced its binding with His_6_-TRPV1 Ct. **b.** Quantification analysis for the GST pull-down experiment. Statistical analysis was performed using unpaired *t*-tests. n= 6, *\*p*< 0.05. **c.** Schematic diagram showing the Gap43-mediated membrane localization experiment. Proteins with the predicted molecular weights were examined by Western blotting. **d.** Left, fluorescence signal observed in HEK 293 cells when TRPV1 Ct-GFP or TRPV1 Ct Δ820-GFP was transfected alone or in combination with Gap43-TRPV1 Ct. Coexpression of Gap43-TRPV1 Ct promoted the membrane localization of TRPV1 Ct-GFP but not of TRPV1 Ct Δ820-GFP. Right, the spatial fluorescence intensity spectrum of GFP.

### The TRPV1Δ820 mutant exhibits reduced current density and increased resistance to trachyphylaxis

To observe the impact of distal Ct deletion on the channel function of TRPV1, we recorded the voltage-induced currents of the TRPV1 channel using the voltage clamp technique. Starting from -100 mV we conducted stepwise increases of voltage by 20 mV until 160 mV **(Fig. 7a)**. In this, both wild-type (WT) TRPV1 and TRPV1 Δ820 mutants could open under voltage stimulation. As the voltage increased, the G/G_max_ also increased. There were no significant differences between the two groups **(Fig. 7b)**, indicating that the TRPV1 Δ820 mutation does not change the voltage dependence properties of TRPV1. This finding was expected as truncation of the distal Ct leaves the extracellular and transmembrane domains unaffected. Consequently, TRPV1 Δ820 displayed a similar activation pattern as WT TRPV1 when subjected to voltage stimulation.

**Fig. 7.**
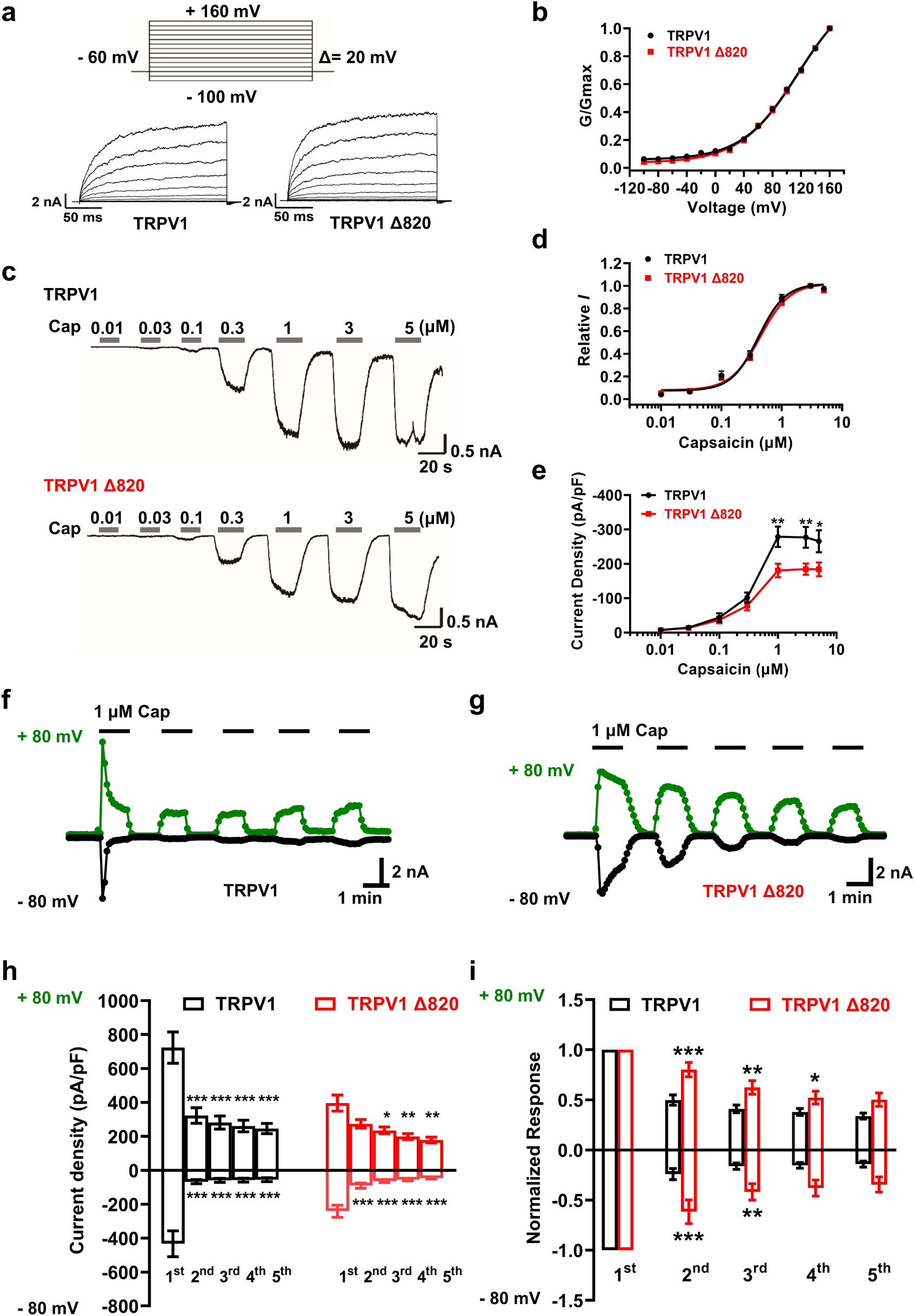
Deleting the distal 19 amino acid residues of TRPV1 Ct decreases the capsaicin-induced current density while increasing channel resistance to tachyphylaxis. **a.** Representative whole-cell current traces from wild type TRPV1 or TRPV1 Δ820 in response to the indicated voltage stimulation in the absence of Ca^2+^. Recordings were conducted starting at -100 mV, increasing stepwise by 20 mV until reaching 160 mV. **b.** The activation curve of TRPV1 and TRPV1 Δ820. **c.** Representative current traces activated by different concentrations of capsaicin in HEK 293 cells overexpressing TRPV1 or TRPV1 Δ820. **d.** Dose-response curves of TRPV1 and TRPV1 Δ820. **e.** Normalized peak current density of TRPV1 and TRPV1 Δ820 by membrane capacitance under the action of different concentrations of capsaicin. For the TRPV1 group, the neuronal number recorded for 0.01, 0.03, 0.1, 0.3, 1, 3, 5 μM capsaicin was 9, 19, 18, 29, 29, 24 and 20, respectively. For the TRPV1 Δ820 group, the corresponding number was 22, 26, 29, 39, 38, 37 and 32, respectively. Statistical analysis was performed using two-way ANOVA. \**p*< 0.05 and ** *p*< 0.01. **f, g.** Sample whole-cell current traces from wild type TRPV1 (**f**) or TRPV1 Δ820 (**g**) in response to consecutive application of capsaicin with the holding potential at -80 mV or +80 mV. Ca^2+^ was included in the bath solution. **h.** Peak current density of TRPV1 and TRPV1 Δ820 normalized by membrane capacitance under the consecutive application of capsaicin (1 μM). For the TRPV1 group, n= 10; For TRPV1 Δ820 group, n= 7. Statistical analysis was performed using Two-way ANOVA followed by the Sidak’s multiple comparisons test. \**p*< 0.05, \*\**p*< 0.01 and \*\*\**p*< 0.001. **i.** Normalized analysis of the data in (h). Statistical analysis was performed using Two-way ANOVA followed by the Sidak’s multiple comparisons test. \**p*< 0.05, \*\**p*< 0.01 and \*\*\**p* < 0.001.

We next recorded the currents of WT TRPV1 and TRPV1 Δ820 under the stimulation of different doses of capsaicin for 20 s **(Fig. 7c)**. Since the experimental system did not contain Ca^2+^, TRPV1 channels did not undergo desensitization and remained highly responsive to capsaicin. In the dose-response curve, there were no differences in the slope and EC_50_ between WT TRPV1 and TRPV1 Δ820 **(Fig. 7d)**, indicating similarity in their capsaicin sensitivity. This finding might be explained that activation of TRPV1 by capsaicin requires two “gates”, with the upper gate located near the pore’s selectivity filter and the lower gate near the S6 crossing^29^. TRPV1 Δ820 did not affect the critical domains for channel opening. We then compared the current density of WT TRPV1 and TRPV1 Δ820 induced by capsaicin. Under low concentrations of capsaicin (0.01-0.3 μM), the current densities of WT TRPV1 and TRPV1 Δ820 were comparable **(Fig. 7e)**. Considering that the transient expression strategy was used in this study, even though the expression level of TRPV1 Δ820 on the cytomembrane was relatively low, the overexpression system guaranteed its high-level expression and ensured an adequate response to low concentrations of capsaicin. However, when the concentration of capsaicin increased to 1 μM, the responsiveness of TRPV1 Δ820 to capsaicin began to decrease significantly compared to WT TRPV1. With increasing concentrations of capsaicin (1 μM, 3 μM, and 5 μM), TRPV1 Δ820 exhibited correspondingly and significantly weaker responsiveness to capsaicin, leading to smaller current densities when compared to WT TRPV1.

In addition, we examined the tachyphylaxis induced by repetitive application of capsaicin (1 μM) in the presence of Ca^2+^ in the recording solution. The TRPV1 current showed a dramatic decline in the first capsaicin application and remained at a low level during all subsequent capsaicin stimulations **(Fig. 7f, h)**. This was considered to be tachyphylaxis of TRPV1 channels. Of note, the maximal current to the first capsaicin application was lowered in TRPV1 Δ820 accompanied with slow inactivation kinetics. Meanwhile, the magnitude of decline in the current density was greatly attenuated in TRPV1 Δ820 **(Fig. 7f, h)**. To compare the tachyphylaxis quantitatively, normalized responses were calculated **(Fig. 7i)**. The data revealed that the magnitude of current decline from the 2^nd^ to 4^th^ application of capsaicin was significantly diminished in the TRPV1 Δ820 group, suggesting TRPV1 Δ820 to be more resistant to capsaicin-induced tachyphylaxis.

In summary, the current work revealed that stimulation of TRPV1 under non-toxic doses of capsaicin triggers calpain-1 activation. This cleaves the Ct at G819/S820 site. The truncated TRPV1 Δ820 thereby shows reduced cell membrane localization and channel current due to increased receptor internalization and impaired subunit assembly. Calpain-1 is concluded as a critical Ca^2+^-dependent effector that follows TRPV1 activation and mediates the feedback regulation of TRPV1 **(Fig. 8)**.

**Fig. 8.**
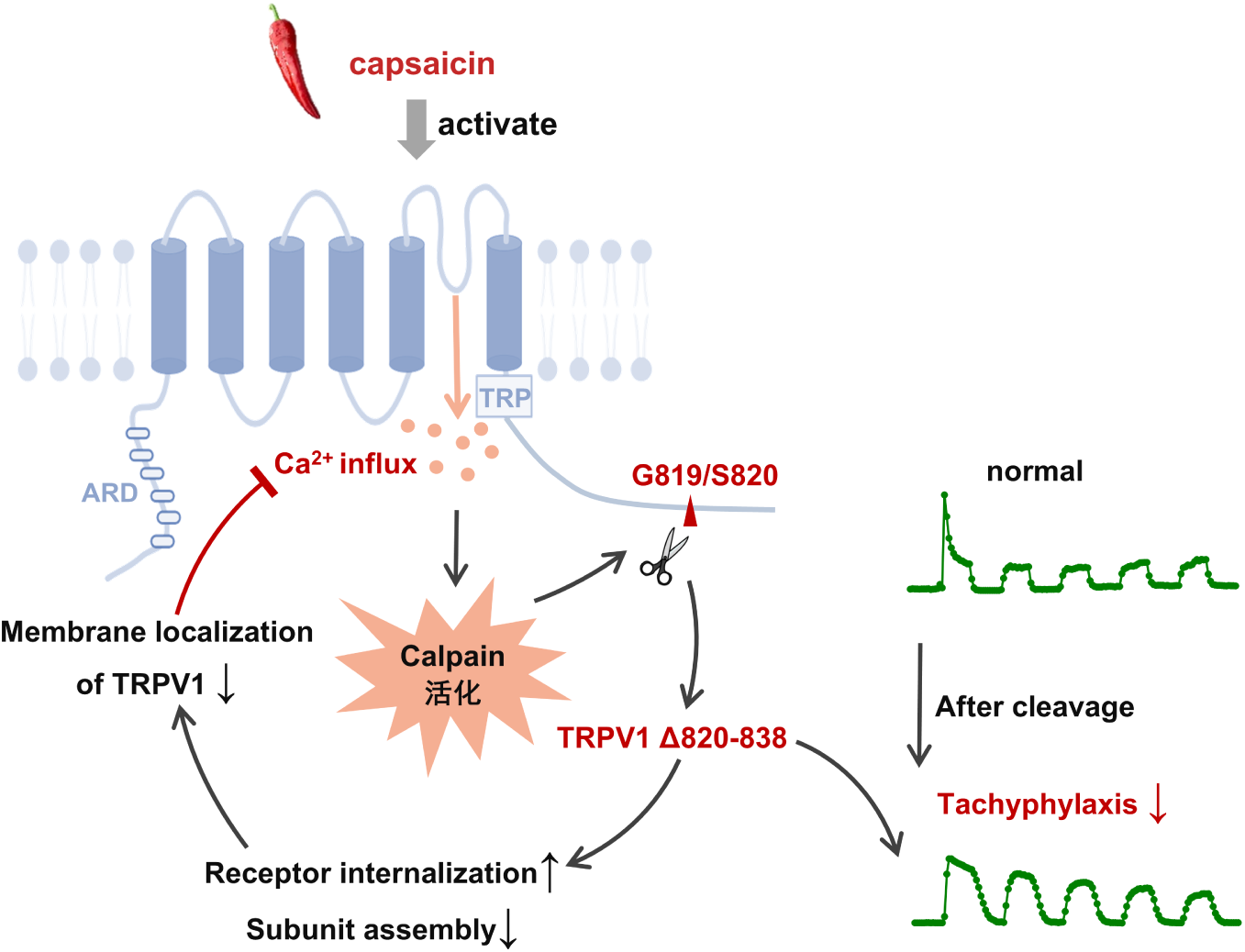
Schematic diagram of the current work, which revealed the direct action of calpain on TRPV1 protein and subsequent regulation of channel function. Calpain-mediated cleavage of TRPV1 Ct plays a finely balanced role in the modulating of TRPV1 function. On one hand, it suppresses the cell membrane localization of TRPV1, which lowers channel function. On the other hand, it renders channel resistance to tachyphylaxis. This results in the channel function beingt maintained within an appropriate range. This dual-regulatory action of calpain might counteract the excessive sensory adaptation brought about by TRPV1 desensitization and protect the body from the potential injury caused by sensory adaptation.

## Discussion

TRPV1 not only plays a key role in thermal perception but is also crucial for heat hyperalgesia and sensitization of peripheral sensory neurons. This makes it a significant target for analgesic drug development. The classic TRPV1 agonist, capsaicin, exerts analgesic effects partially by inducing TRPV1 desensitization, a process that includes both acute desensitization and tachyphylaxis^30^. Ca^2+^-dependent proteins such as calcineurin, PLC, and CaM have been shown to contribute to TRPV1 desensitization^13, 14, 22, 31, 32, 33^. Given the Ca^2+^-dependence of TRPV1 desensitization and the reduced protein level of TRPV1 after capsaicin treatment, we explored the potential role of calpain, a Ca^2+^-dependent cysteine protease, in TRPV1 desensitization. The main findings in this work include: (1) Activation of TRPV1 by a non-toxic dose of capsaicin is accompanied by the activation of calpain. (2) Calpain-1 cleaves distal Ct of TRPV1 at the G819/S820 site. (3) Calpain promotes TRPV1 internalization with truncated TRPV1 showing reduced subunit assembly. (4) Truncated TRPV1 exhibits reduced current density, but attenuated channel tachyphylaxis. Accordingly, our study reveals the elegant regulatory role of calpain-1 in modulating TRPV1. On one hand, calpain-1 reduces TRPV1 current density by decreasing the channel’s localization at the plasma membrane; on the other hand, it protects the channel against tachyphylaxis. This combines to ensure TRPV1 activity is maintained within an appropriate range.

## TRPV1-Ca^2+^-calpain pathway

Previous studies have suggested the existence of a TRPV1-Ca^2+^-calpain pathway as usually associated with cell injury or cell death. For example, in a diabetic rat model, activation of TRPV1 in the large-diameter dorsal root ganglion (DRG) neurons caused cellular stress and injury involving the activation of caspase and calpain. However, this effect was not observed in DRG neurons from healthy rats^34^. This finding is corroborated in the current work. Here, treatment with capsaicin (1 μM, 10 minutes) involved no accompanying activation of caspase 3 **(Fig. 1c)**. In intestinal cells, epidermal growth factor receptor (EGFR) signaling was coupled with activation of TRPV1, which was followed by calpain activation and the cleavage of protein tyrosine phosphatase 1B (PTP1B). Consequently, PTP1B activity was increased and tumorigenesis was suppressed through dephosphorylation of EGFR^35^. In macrophages, excessive nitric oxide activated TRPV1, which in turn triggered calpain-mediated cleavage of liver X receptor α (LXRα), ultimately impairing cholesterol efflux^36^. Additionally, activation of the TRPV1-Ca^2+^-calpain-2 pathway in outer hair cells was associated with cisplatin-induced oxidative stress^37^. Recent studies have indicated that axonal degeneration followed by high-dose capsaicin (1 mM) depends on the action of calpain^38, 39^. Combined with the fact that a large-dose of capsaicin has been used to ablate DRG neurons or achieve anti-tumor effects^40^, widespread activation of the TRPV1-Ca^2+^-calpain pathway, particularly the activation of calpain-2, is noted as detrimental to cell survival.

In the current work, we demonstrated that TRPV1-Ca^2+^-calpain pathway activated by a regular dose (1 μM) of capsaicin is not associated with cell toxic injury. This may be due to a region-specific activation pattern of calpain, which starts from the region under the plasma membrane and then diffuses throughout the whole cell. This is distinct from the widespread activation of calpain following a large dose of capsaicin. In addition, the direct contribution of calpain-1 to the cleavage of TRPV1 was demonstrated in the current studies, although we could not completely exclude the role of calpain-2. Calpain-1 and calpain-2, the two major calpain isoforms, have actually been shown to execute opposite functions in synaptic plasticity and neuroprotection^41, 42, 43^. Furthermore, our work extended previous findings by revealing a closed regulatory loop of TRPV1-Ca^2+^-calpain-TRPV1. Calpain, activated by Ca^2+^ influx through TRPV1 channels, acts on the intracellular segments of TRPV1 conversely, which forms a feedback regulatory loop. The functional significance of this regulatory loop may operate in the maintenance of TRPV1 activity within an appropriate range.

### Characteristics of calpain-mediated substrate cleavage

A large number of substrates for calpain have been identified. These include cytoskeletal proteins, receptors and ion channels, enzymes, kinases and phosphatases. Notably, substrate recognition of calpain is independent of certain amino acid residues or consensus sequences, but depends on the overall three-dimensional structure of the target protein^27^. This functional property of calpain might lead to the failure of our attempts to identify the critical amino acid residues involved in the cleavage of the TRPV1 Ct by calpain. It has been reported that protease-mediated cleavage of substrates tends to occur among three types of secondary structure, namely extended loops, α-helices and β-sheets^44, 45^, all of which are detectable in the protein sequence of the TRPV1 Ct^5, 46^. However, further studies are required to precisely determine the secondary structure or amino acid residues involved in the action of calpain on TRPV1 Ct.

Another feature of calpain’s action is that it hydrolyzes substrates to a limited extent, thereby conferring upon it the role of a modulator protease. For example, calpain-mediated cleavage of Ca_v_1.1 or Ca_v_1.2 produces an autoinhibitory fragment in the distal Ct. This then inhibits channel function through interaction with the proximal Ct^47, 48^. Conversely, the truncated transient receptor potential canonical 5 (TRPC5), as produced by the action of calpain and which occurs in the TRPC5 Ct, exhibits a potentiated channel function^49^. Analogically, calpain-mediated cleavage of TRPV1 also occurs at the distal Ct. Truncated TRPV1 therefore exhibits a finely balanced change in channel function. However, the possible functions of small fragment 820-838 amino acid residues remain to be further determined.

### Functional regulation of TRPV1 resulted from calpain-mediated cleavage of its C terminus

The action of calpain on TRPV1 leads to the generation of truncated TRPV1 lacking a distal Ct. The TRPV1 Ct has been shown to play complex regulatory roles in TRPV1. For example, the TRP domain located in the proximal Ct of TRPV1 plays a critical role in subunit assembly and channel activation^50, 51^. The TRP domain, particularly K710, is involved in the activation or potentiation of TRPV1 by lysophosphatidic acid and PIP_252, 53_. Distal to this PIP_2_ binding site, there is another distal PIP_2_ binding site (777-820 amino acid residues), which exhibits an inhibitory effect on TRPV1^54, 55^. Overlapping with this segment lies the sequence of 767-801 amino acid residues, where particularly L796 and W787 act as the C terminal binding sites for CaM on TRPV1^14, 16^. As both PIP_2_ and CaM contribute to TRPV1 desensitization, whether there is functional relationship between the distal TRPV1 Ct and the other regions of TRPV1 Ct remains to be determined. On the other hand, recent studies have indicated that TRPV1 desensitization is accompanied with Nt-Ct interactions. The truncation of distal TRPV1 Ct by calpain might destroy this interaction, leading to attenuated receptor desensitization.

In addition to the possible effects on protein-protein interaction, truncation at G819/S820 leads to the loss of post-translational modification sites of TRPV1. For example, K822 is a critical site for the small ubiquitin-like modifier (SUMO) modification of TRPV1, which specifically enhances the channel’s sensitivity to heat, but not capsaicin, protons or voltage^56^. TRPV1 K822 SUMOylation is essential for the development of thermal hyperalgesia during inflammatory pain and the suppression of itching sensation via interrupting the binding between TRPV1 and histamine receptor type 1 (H1R)^57^. Therefore, calpain-mediated cleavage of the TRPV1 Ct likely reduces TRPV1 sensitization to heat stimulation by abolishing its SUMOylation. Additionally, S774 and S820 are the phosphorylation sites located in the C terminal region of TRPV1 by protein kinase A (PKA), which promotes TRPV1 sensitization and reduces desensitization in a phosphorylation-dependent manner^13, 58^. Despite this, the contribution of these sites to the regulatory action of PKA on TRPV1 has not been acknowledged and the exact role of S820 in the functional regulation of TRPV1 remains to be determined.

In summary, the current studies reveal the dual regulatory role of calpain on TRPV1, where both reduction of the TRPV1 channel current and concurrent attenuation of receptor desensitization is achieved. Overall, the function of TRPV1 is downregulated after calpain activation, ensuring that TRPV1 channel activity remains within a narrow, appropriate range. However, there are some limitations for the current studies. For example, whether activation of TRPV1 couples with the elevated activity of calpain and subsequent cleavage of TRPV1 Ct *in vivo* remains unknown. Future measurement in the primary cultured sensory neurons or DRG tissue are also required. In addition, the question of how we can develop ways of specifically promoting the action of calpain on TRPV1 without affecting the other substrates of calpain remains a critical issue. Addressing these matters could lead to the development of novel analgesic drugs targeting TRPV1 towards improved clinical treatment for chronic pain and reduction of global disease burden.

## Materials and Methods

### Plasmid construction

PCR primers were designed using SnapGene and were synthesized by RuiBiotech (Beijing, China). Detailed information is shown in **Table 1**. The construction approach of eukaryotic expression plasmids of GST-TRPV1 and TRPV1-GFP, the prokaryotic expression plasmids of GST-TRPV1 Nt and GST-TRPV1 Ct was described in our previous studies^59, 60^. PCR products were purified using a TIANgel Purification kit (TIANGEN, DP219-02). DNA concentrations were measured using a NanoDrop spectrophotometer. Then a seamless ligation system (GenStar, T196) was prepared at a 3:1 ratio (insert fragment: vector) and incubated at 50°C for 15 minutes. For vectors, pEGFP-N1 was used for the eukaryotic expression and PGEX-5X-1 and pET-28a(+)were used for the prokaryotic expression. The reaction products were then transformed into competent cells (DH5α for eukaryotic expression, BL21 for prokaryotic expression). All plasmids were verified by DNA sequencing.

### Cell culture, plasmid transfection and drug treatment

The HEK 293 cell line was obtained from the Cell Resource Center, Peking Union Medical College. CHO cells stably expressing TRPV1 were as described in our previous studies^59, 61^. The culture medium was DMEM (MACGENE) and Ham’s F-12 (MACGENE), both individually supplemented with 10% FBS (Gibco). Cells were cultured in a humidified atmosphere of 95% air and 5% CO_2_ at 37°C.

Cells were passaged 24 hours before transfection to achieve 60-80% confluency. Transfection was performed using Lipofectamine 3000 (Invitrogen). 1 μg empty vector or 2 μg recombinant plasmids were transfected in a 35 mm dish unless otherwise specified. Plasmid DNA and P3000 reagent (1:2 ratio) were mixed in 100 μl of Opti-MEM and incubated for 5 minutes. Meanwhile, 100 μl lipofectamine 3000 was diluted with equal volume of Opti-MEM and the mixture was added to the previous DNA mixture. After 15 minutes, 200 μl of the transfection mix was added to cells in 800 μl of fresh medium. Cells were harvested 24-48 hours post-transfection.

Capsaicin (Sigma, 404-86-4) was dissolved in alcohol to prepare a stock solution of 100 mM, which was diluted to the appropriate concentration just before use. For the inhibitors experiment, Calpeptin (Merck, 03-34-0051), MDL28170 (Merck, M6690), Z-VAD-FMK (Merck, V116), or MG132 (Merck, M8699) was added 30 minutes in advance.

### Cell staining and capture of fluorescence images

ER-Tracker Red (Yeason, 40764ES20) was diluted to 0.2 μM in Hank’s balanced salt solution (Yeason, 60147ES76) and added to the culture dish. The cells were then incubated at 37°C for 20 min and subsequently fixed with 4% paraformaldehyde and rinsed with PBS. Cellular nuclei were stained with Hoechst 33258 dye (Yeasen, 40730ES03) for 10 minutes at room temperature. After rinsing with PBS, cells were mounted using a mounting medium (ZSGB-BIO, Zli-9556). Images were captured using an Olympus FV 3000 confocal laser scanning microscope with a 100× HC PL APO objective (NA 1.45; oil). Fluorescence intensity was quantified using CellSens (Olympus).

### Calpain activity assay

Calpain activity was measured using the calpain activity assay kit (Abcam, ab65308) following the manufacturer’s instructions. Cells were lysed with extraction buffer on ice for 20 minutes, then scraped. Cellular lysates were centrifuged at 10,000 g for 2 minutes. The supernatant was collected and protein concentration was determined using a BCA assay kit (Pierce). 100-200 μg protein was subsequently isolated for incubation with reaction buffer at 37°C for 1 hour. Fluorescence intensity was then measured at 400 nm excitation and 505 nm emission using a spectrophotometer.

### Prokaryotic protein expression and purification

The fresh bacterial culture was incubated at 37°C, 250 rpm for 3-4 hours, 0.1 mM IPTG (Genview) added, and incubation continued at 20°C, 250 rpm overnight. The next day bacteria were centrifuged at 4°C, 4,000 rpm for 10-15 minutes with the pellet then collected and lysed with the appropriate lysis buffer. Triton X-100 was added and the lysates were sonicated on ice. After centrifugation at 4°C, 13,000 rpm for 30 minutes, the supernatant was collected.

Resin beads (Yeason, 20507ES10 or Qiagen, 30210) were loaded into a chromatography column (Smart-life, SLM007). The column was washed 10 times using the column volume of water and a further 5 times using the column volume of washing buffer. The supernatant was then added at 1 ml/min and the column washed again with 10-15 times the column volume of washing buffer. After that, 5-10 times column volume of elution buffer was added at 1 ml/minute and the eluate collected and concentrated using microcentrifuge tubes (Millipore, UFC505008). The protein concentration was then measured and the protein aliquoted. The aliquots were stored at - 80°C until use.

### GST-pull down assay

The eluate of His_6_-tagged proteins was added to the glutathione-Sepharose resin with bound GST or GST-tag proteins. The mixture was suspended at 4°C for 2 hours. Six column volumes of 0.1% Triton X-100/TBS were added and the mixture was centrifuged at 2,000 rpm for 2 minutes at 4°C. This procedure was repeated 6 times to remove unspecific binding. Then 10-20 μl of elution buffer was added and the mixture was suspended at 4°C for 1 hour. Thereafter, supernatant was collected and 5× SDS-PAGE loading buffer was added followed by SDS-PAGE electrophoresis.

### Western Blotting

Approximately 50 μg protein was loaded into each lane onto 10% polyacrylamide gels and electrophoresed at 80-120 V. The proteins were then transferred to a nitrocellulose membrane (Pall) using a wet blot transfer system (Bio-Rad Laboratories) at 200 mA for 1-2 hours and the membranes were blocked with 5% non-fat milk in TBST (tris-buffered saline with 0.05% Tween 20) for 1 hour. Further incubation with primary antibodies (anti-GST antibody, Applygen, C1303, 1:3000; anti-His_6_ antibody, Applygen, C1301, 1:3000; anti-Flag antibody, Applygen, C1305, 1:1000; anti-GFP antibody, Abcam, ab290, 1:1000; anti-TRPV1 Ct antibody, Alomone, ACC030, 1:1000; anti-TRPV1 extracellular domain antibdoy, Alomone, ACC029, 1:500; anti-transferrin antibody, Abcam, 1:1000; anti-β-actin antibody, ZSGB-BIO, TA-09, 1:2000) was conducted overnight at 4°C. After washing with TBST for 3 times, the membranes were incubated with the appropriate horseradish peroxidase-conjugated secondary antibody (Jackson ImmunoReseach) for 1 hour at room temperature. Signals were detected using the chemiluminescent HRP substrate kit (Millipore).

### Protein enrichment by immunoprecipitation

rProtein A/G agarose resin (Yeason, 36403ES05) was pre-cleaned and PBS and anti-GFP antibody (1:100) diluted with TBS added to the resin and suspended at 4°C for over 3 hours. Cellular lysates were then added and the mixture was suspended at 4°C overnight. The resulting resin was washed six times with 0.1% Triton X-100/TBS. Elution buffer was added and the mixture suspended at room temperature for 10 min. Finally, the mixture was centrifuged at 13,000 rpm for 10 minutes at 4°C and the supernatant was transferred to a clean tube.

### Protein treatment by calpain *in vitro*

Protein substrate was mixed with equal volume of 2× calpain digestion solution (80 mM Hepes, 10 mM DTT, pH 7.2). Appropriate volumes of calpain-1 (Merck, 208712) were added. Thereafter, CaCl_2_ was added to a final concentration of 1 mM. The mixture was incubated at 37°C for the required time.

### Biotinylation of the cell surface protein and assay of receptor internalization

Cell cultures were washed twice with ice-cold DPBS and incubated for 30 minutes at 4°C with EZ-Link Sulfo-NHS-LC-Biotin (Thermo Scientific, 1 mg/ml) to biotinylate the cell surface proteins. Excess biotin reagent was quenched by washing the cells with PBS containing 100 mM glycine. Thereafter, cells were lysed and the lysates were centrifuged at 12,000 g for 5 minutes to yield protein extracts in the supernatant. The protein extracts were then incubated with streptavidin agarose resin 6FF (Yeason, 20512ES08) for 2 hours at 4°C to capture biotinylated surface proteins. After being washed with lysis buffer, proteins were eluted by boiling for 5 minutes with 5× SDS-PAGE loading buffer and the sample was subjected to SDS-PAGE electrophoresis.

For the experiment of receptor internalization, cells were incubated at 37°C for 10 minutes after the biotinylation labeling. Biotin that remained at the cell surface was then removed by incubation with 100 mM MESNA solution and cells washed with DPBS, lysed with the lysis buffer, for final SDS-PAGE electrophoresis.

### FRET experiment

FRET was performed on a Leica TCS SP8 confocal microscope with a 63× HC PL APO objective (NA 1.40, oil) using FRET AB (acceptor bleaching) Leica software modules. Cells were plated at low density on a confocal dish and transfected with the pCMV-Calpainsensor (Addgene, 36182) or pEYFP-N1 (YouBio, VT1105) and pECF-N1 (YouBio, VT1113) plasmids. For FRET AB, regions of interest were drawn and excited with the 514 nm laser line to bleach the YFP channel. Both a prebleach image and a postbleach image of the CFP and YFP channels were acquired, and the intensity change of CFP was analysed using the Leica FRET AB Wizard software (LAS AF). The FRET efficiency was calculated using the following formula: FRET% =((CFPAP-CFPBP)/CFPAP) × 100 %.

### Mass spectrometry analysis

The bands were excised from the SDS-PAGE gel, then reduced with 10 mM DTT and alkylated with 55 mM iodoacetamide. Then in-gel digestion was carried out with trypsin (Promega) in 25 mM ammonium bicarbonate at 37°C overnight. The digested sample was loaded onto a C18 trap and followed by nano-LC-ESI-MS/MS analysis. The peptides were sequentially eluted from the HPLC column with a gradient of 2-45% of Buffer B (acetonitrile:water:acetic acid, 80:19.9:0.1) in Buffer A (acetonitrile:water:acetic acid, 0:99.9:0.1) at a flow rate of 300 nl/min. The eluted peptides were sprayed directly from the tip of the capillary column to the mass spectrometer (Thermo OE480), which was operated in the data-dependent mode. The initial MS scan recorded the mass to charge (m/z) ratios of ions over the mass range from 450-1800 Da. The 20 most abundant ions were automatically selected for subsequent collision-activated dissociation.

All MS/MS data were searched against a rat protein database downloaded from the uniprot database using Proteome Discoverer (Thermo). All searches were performed using a precursor mass tolerance of 0.02 Da calculated using average isotopic masses. Variable modification was set for methionine with the addition of 15.999 Da to represent methionine oxidation, static modification was set for cysteine with the addition of 57.052 Da to represent cysteine carboxyamidation. A fragment ion mass tolerance of 20 ppm was used. Enzyme cleavage specificity was set to trypsin and no more than two missed cleavages were permitted. The SEQUEST outputs were then analyzed.

### Patch clamp recordings

HEK 293 cells were cultured in DMEM supplemented with 10% FBS, 50 units/ml penicillin, and 50 μg/ml streptomycin. When they reached 70-80% confluence the cells were transfected with the desired DNA constructs using Lipofectamine 3000 (Invitrogen) following the manufacturer’s protocol. After 18-24 hours of transfection, the cells were trypsinized and plated onto glass coverslips. Measurements were taken 6-12 hours after plating.

Patch clamp recordings were conducted using voltage-clamped on a HEKA EPC10 amplifier with PatchMaster software (HEKA) in the conventional whole-cell configuration. Patch pipettes were pulled from borosilicate glass to a resistance of 2-4 MΩ. The pipette solution contained 140 mM CsCl, 5 mM EGTA, and 10 mM HEPES. The pH was adjusted to 7.2 using CsOH. The bath solution for recording consisted of 145 mM NaCl, 5 mM KCl, 1 mM MgCl_2_, 1 mM EGTA, 10 mM glucose, and 10 mM HEPES. The pH was adjusted to 7.4 with NaOH. For desensitization testing, the bath solution containing physiological 1.8 mM Ca^2+^ contained: 145 mM NaCl, 5 mM KCl, 1 mM MgCl_2_, 1.8 mM CaCl_2_, 10 mM glucose, and 10 mM HEPES. For voltage-dependence experiments, a voltage step protocol was used. This protocol consisted of 200-ms depolarizing pulses ranging from −100 mV to 160 mV at 20-mV increments. The protocol was triggered from the holding potential of -60 mV. When recording, serial resistance was compensated by 60% to reduce voltage errors. For desensitization recording, repeatedly stimulated with 1 µM capsaicin for 1 min followed by 1-min washouts at a holding potential of -60 mV or voltage ramps (1500 ms) from -100 mV to +100 mV were applied every 5 s from a holding potential of 0 mV. The current signal was sampled at 5 kHz and filtered at 2.9 kHz. All measurements were performed at room temperature (22–24°C). To prepare capsaicin for the experiments, it was dissolved in pure ethanol to create a stock solution. The solution was then diluted in the bath solution to the desired concentration immediately before use.

During recording, a rapid solution changer with a gravity-driven perfusion system (RSC-200, Bio-Logic) was used to apply the capsaicin solutions. Each solution was delivered through a separate tube to prevent mixing. The G-V curves were generated by converting the maximum current values to conductance using the formula G = I / (V - VR), where G represents the conductance, I represents the peak current, V represents the command pulse potential, and VR represents the reversal potential of the ionic current obtained from the I-V curves. Normalized conductance (G/G_max_) was obtained from the steady-state current at the end of each pulse and fitted to the Boltzmann equation: G/G_max_ = 1/(1 + exp[-(Vm - V ½)/k]).

### Statistical analysis

Data were shown as means ± SEM and analyzed using GraphPad Prism (GraphPad Software). Differences between two groups were compared using unpaired *t*-tests with or without Welch’s correction. Differences between multiple groups were compared using One-way ANOVA followed by the Dunnett’s multiple comparisons test, or Two-way ANOVA followed by the Sidak’s multiple comparisons test. A *p* value < 0.05 was considered statistically significant.

### Reporting summary

Further information on research design is available in the Nature Portfolio Reporting Summary linked to this article.

## Data availability

The data that supports this study are available from the corresponding authors upon request.

## Acknowledgement

We would like to thank Dr Xiaomeng Shi in State Key Laboratory of Natural and Biomimetic Drugs of Peking University for her help in the experiment of mass spectrometry.

## Author contributions

Y.Z., P.L. and Y.W. were involved in the overall design of the studies. J.Y.J., Y.Z., J.Y. and S.W.G. performed the biochemical experiments. J.Y.J., J.X.X., Y.Z. and J.L. performed the fluorescence imaging experiments. X.C.W. completed the patch clamp recording. Y.Z., J.Y.J. and X.C.W. wrote the first draft. P.L., Y. Z. and Y.W. revised the manuscript.

