## Supplementary Figures for "Calpain cleaves the carboxyl terminus of TRPV1 and modulates receptor tachyphylaxis"

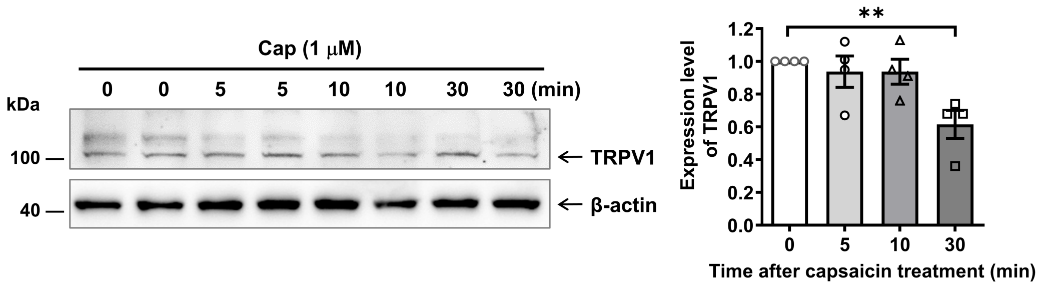


**Supplementary Fig. 1. Capsaicin treatment did not induce a significant decrease of TRPV1 protein in CHO-TRPV1 cells until 30 minutes after treatment.** Statistical analysis was performed using One-way ANOVA followed by the Dunnett's multiple comparisons test, n= 4, $^{**}p < 0.01$.

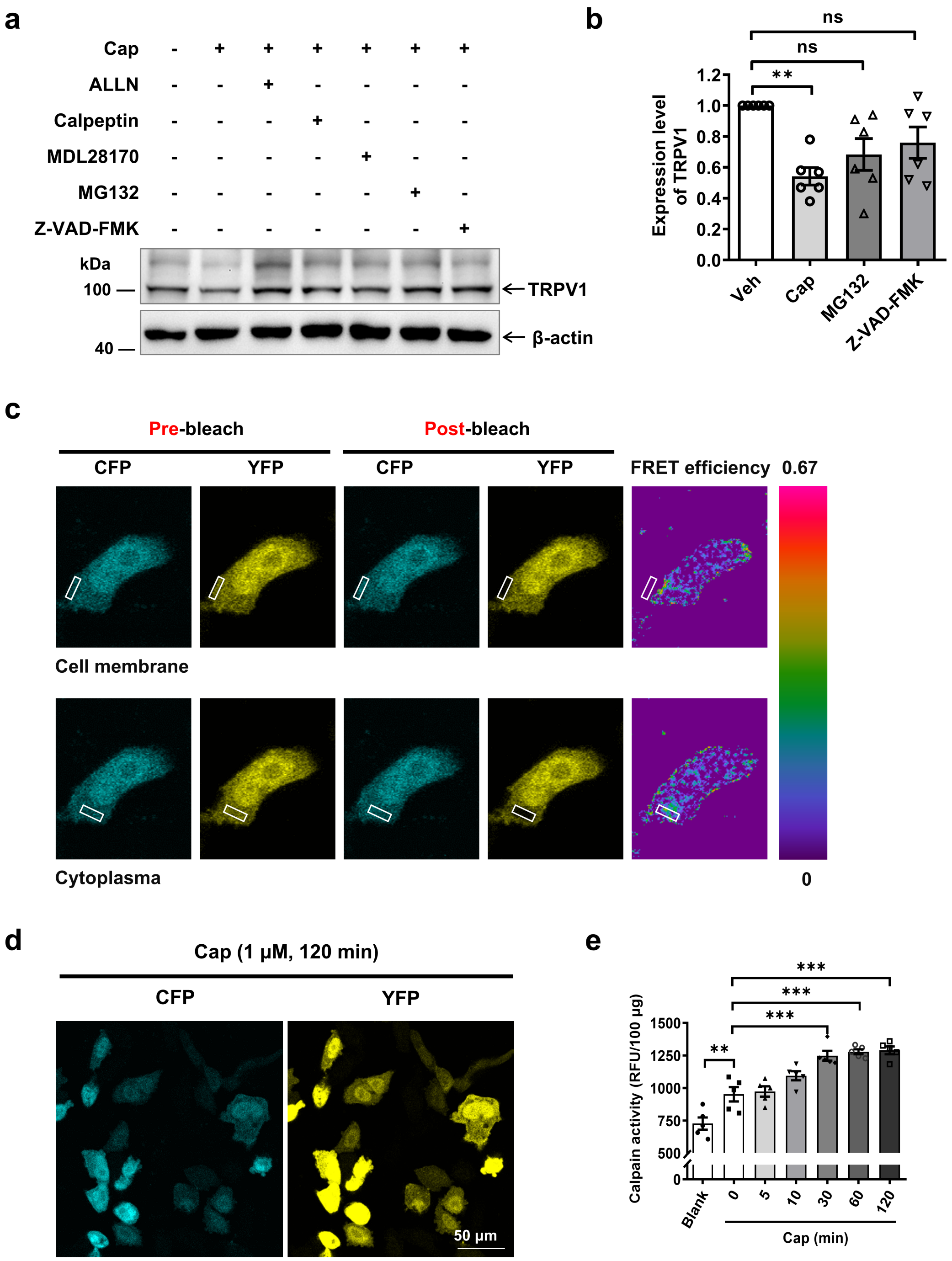


**Supplementary Fig. 2. Activation of calpain following capsaicin stimulation of TRPV1. a, b.** Pretreatment with the inhibitor of ubiquitin, MG132 (25 μM), or the caspase inhibitor, Z-VAD-FMK (10 μM), failed to block the activity-dependent reduction of TRPV1 protein. Statistical analysis was performed using one-way ANOVA followed by the Dunnett's multiple comparisons test, n= 6, $^{**}p< 0.01$. **c.** Representative images of pre-bleaching, post-bleaching and FRET efficiency in the FRET assay. **d.** Representative images of CHO-TRPV1 cells 2 hours post capsaicin stimulation. **e.**

Measurement using a calpain activity assay kit showed that stimulation with capsaicin (1 μM) for 30 minutes, 1 hour, or 2 hours, in CHO-TRPV1 cells increased calpain activity. Statistical analysis was performed using One-way ANOVA followed by the Dunnett's multiple comparisons test, n= 5, $^{**}p < 0.01$ and $^{***}p < 0.001$.

**a**

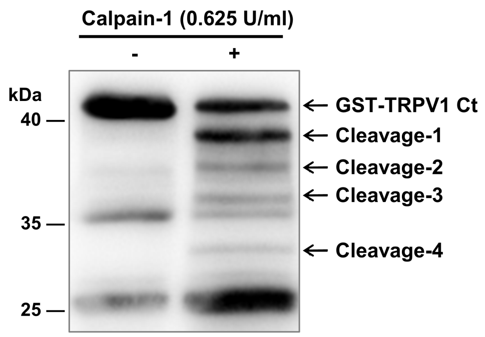


**b**

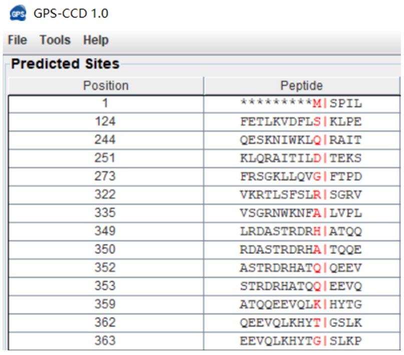

GPS-CCD 1.0

File Tools Help

Predicted Sites

| Position | Peptide |
|---|---|
| 1 | *********M\|SPIL |
| 124 | FETLKVDFLS\|KLPE |
| 244 | QESKNIWKLQ\|RAIT |
| 251 | KLQRAITILD\|TEKS |
| 273 | FRSGKLLQVG\|FTPD |
| 322 | VKRTLSFSLR\|SGRV |
| 335 | VSGRNWKNFA\|LVPL |
| 349 | LRDASTRDRH\|ATQQ |
| 350 | RDASTRDRHA\|TQQE |
| 352 | ASTRDRHATQ\|QEEV |
| 353 | STRDRHATQQ\|EEVQ |
| 359 | ATQQEEVQLK\|HYTG |
| 362 | QEEVQLKHYT\|GSLK |
| 363 | EEVQLKHYTG\|SLKP |

**c** GST-TRPV1-Ct

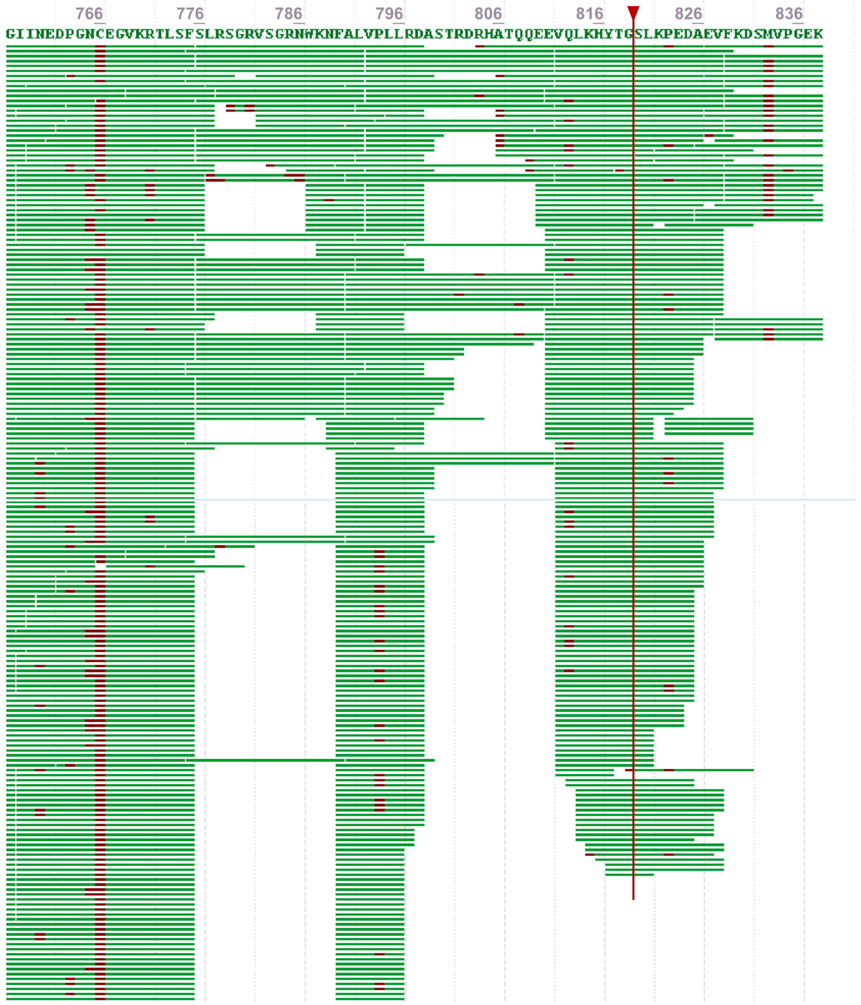

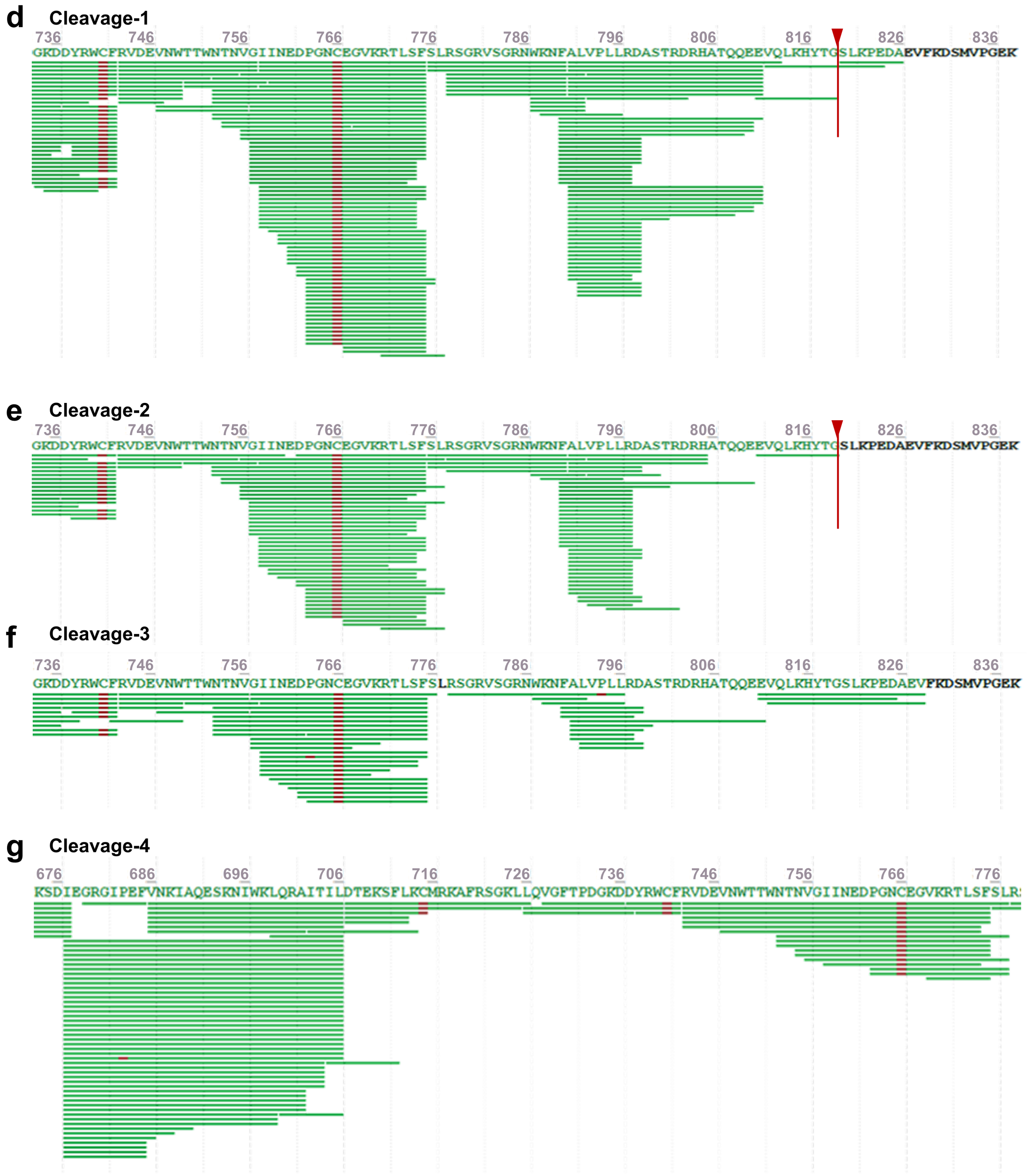


**Supplementary Fig. 3. Mass spectrometry analysis of the cleavage site of TRPV1 Ct by calpain-1. a.** Coomassie brilliant blue staining of GST-TRPV1 Ct protein after incubation with calpain-1. **b.** Calpain cleavage sites predicted by the GPS-CCD program. **c-g.** Mass spectrometry analysis of GST-TRPV1 Ct (**c**) and their associated cleavage fragments, Cleavage-1 to -4, respectively (**d-g**). The upper red arrows denote the potential cleavage sites of calpain.

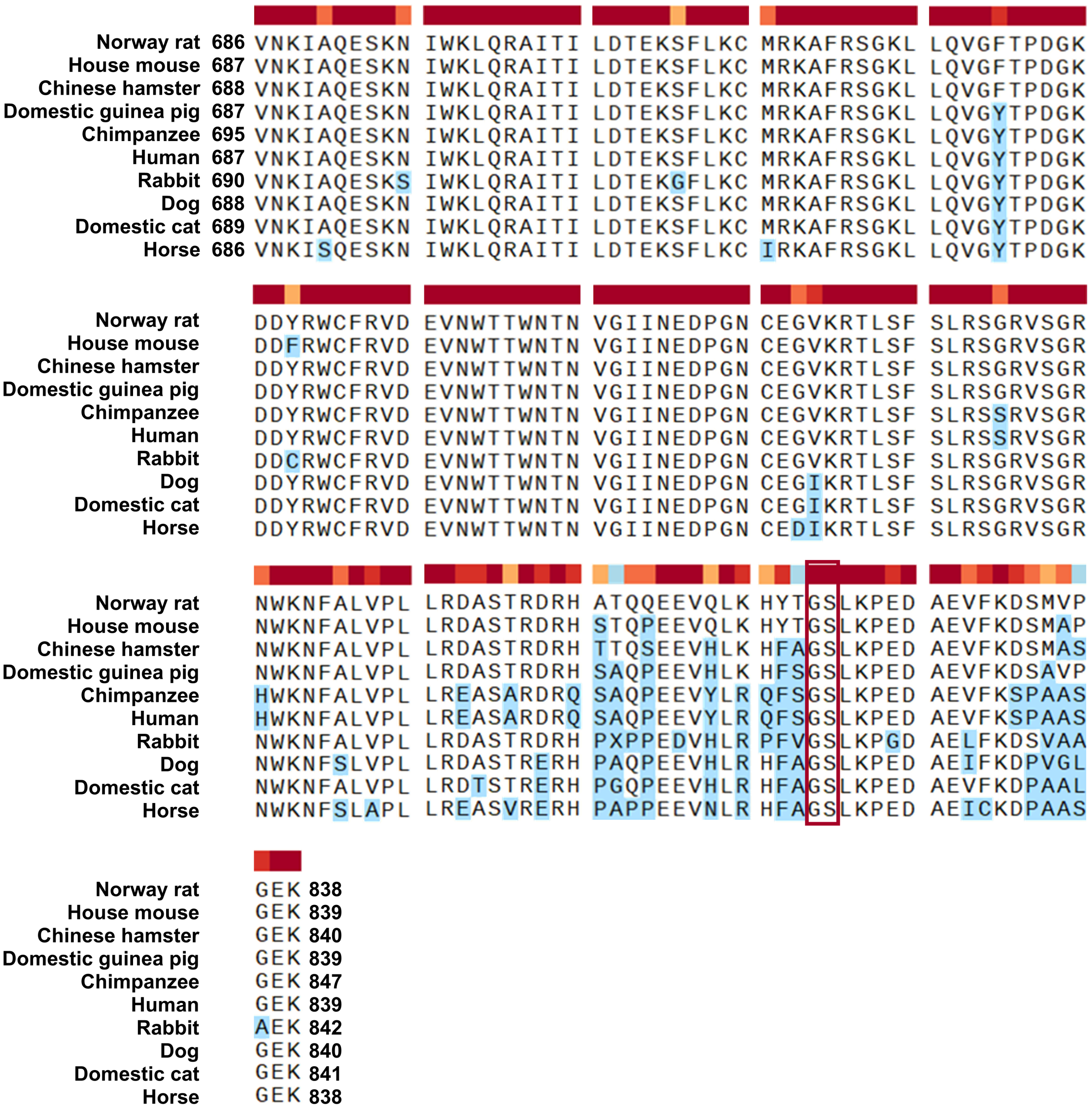


**Supplementary Fig. 4. Alignment of TRPV1 C terminal amino acid sequences from 10 mammals.** The color bar in the top denotes the degree of conservation of amino acid residues. The deeper the color, the higher the conservation. The amino acid residues differing to that of the Norway rat are highlighted in blue.

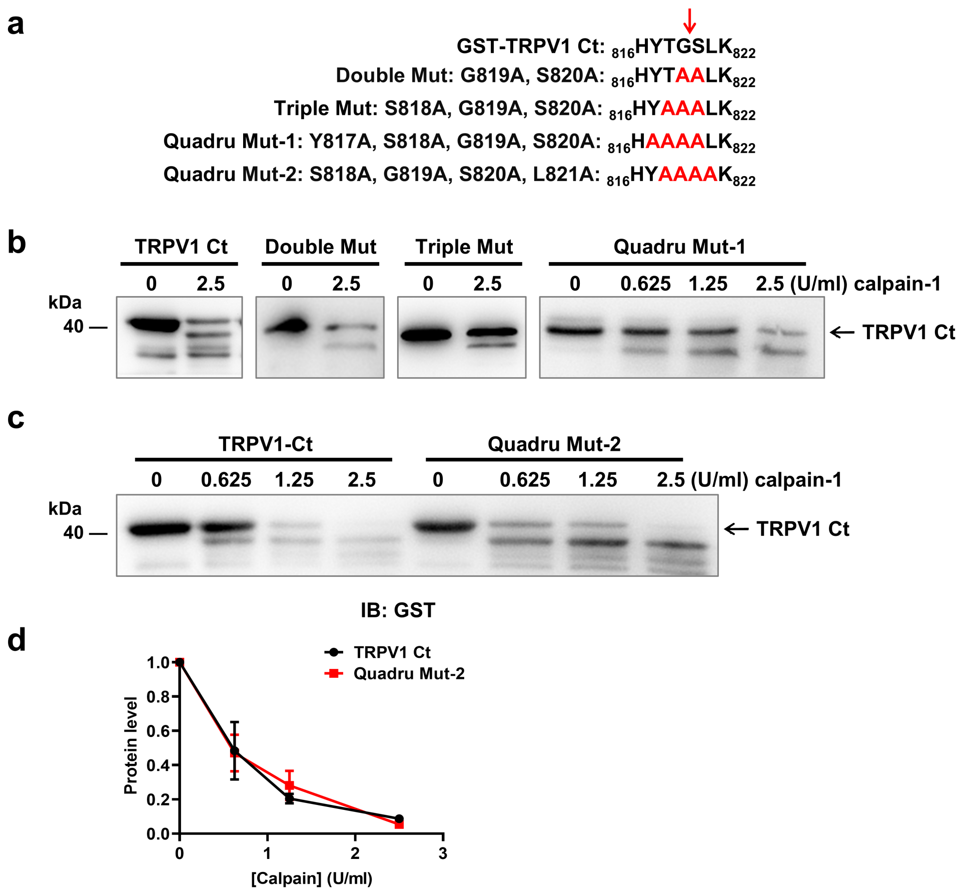


**Supplementary Fig. 5. Mutational analysis for the critical amino acid residues involved in calpain-mediated cleavage of TRPV1 Ct. a.** Construction of the double, triple, and quadruple mutants of GST-TRPV1 Ct around G819/S820 sites. **b.** Effects of the double, triple, or quadruple mutation around G819/S820 sites of TRPV1 Ct on calpain-mediated cleavage of GST-TRPV1 Ct. **c.** Effects of a quadruple mutation around G819/S820 sites on the concentration-dependent cleavage of GST-TRPV1 Ct by calpain. **d.** Quantification analysis of the protein level of GST-TRPV1 Ct or its quadruple mutant 2 (S818A, G819A, S820A, L821A) after incubation with different concentrations of calpain for 15 minutes. The data were analyzed using Two-way ANOVA analysis.

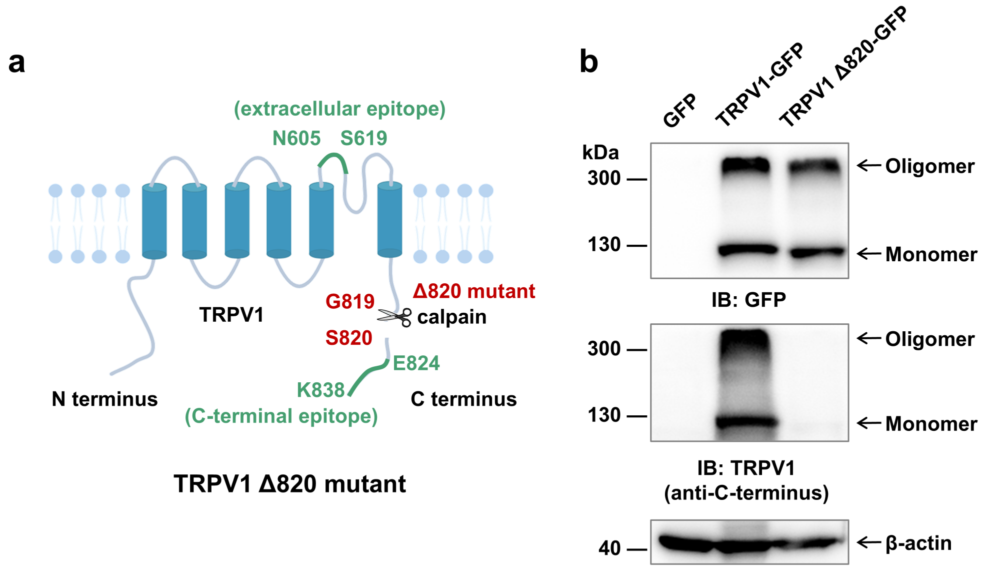


**Supplementary Fig. 6. Verification of TRPV1 Δ820-GFP plasmid. a.** Construction diagram of TRPV1 Δ820-GFP plasmid. **b.** Examination of the TRPV1 Δ820-GFP protein expression using Western blotting.

**Supplementary Table 1. Primers used in the plasmid construction.**

| Recombinant plasmids | Primer | 5’→3’ |
|---|---|---|
| TRPV1 Δ820-GFP | pEGFP-N1-F | ATACGGGAAGGTCGACGGTACCGCGG |
| | pEGFP-N1-R | TTGTTCCATGAGCGCTAGCGGATCTGACG |
| | TRPV1-F | GTCAGATCCGCTAGCGCTCATGGAACAACGGGCTAGCTTAGAC |
| | TRPV1-R | CCGCGGTACCGTCGACCTTCCCGTATAATGCTTCAGTTGAACTTCT |
| TRPV1 Ct-GFP | pEGFP-N1-F | AGGGGAGAAAAGCTTCGAATTCTGCAGTCGAC |
| | pEGFP-N1-R | TCTTGTTGACTAGCGCTAGCGGATCTGACG |
| | TRPV1-F | GCTAGCGCTAGTCAACAAGATTGCACAAGAGAGCAAGA |
| | TRPV1-R | ATTCGAAGCTTTTCTCCCCTGGGACCATGGA |
| TRPV1 Ct Δ820-GFP | pEGFP-N1-F | TTATACGGGAGTACCGCGGGCCCG |
| | pEGFP-N1-R | TCTTGTTGACAATTCGAAGCTTGAGCTCGAGATCT |
| | TRPV1-F | GCTTCGAATTGTCAACAAGATTGCACAAGAGAGCAAGA |
| | TRPV1-R | CCCGCGGTACTCCCGTATAATGCTTCAGTTGAACTTCTTCC |

| | | |
|---|---|---|
| GAP43-TRPV1 Ct | pcDNA3.1-GAP43-F | AGGGGAGAAAAATTCTGCAGATATCCAGCACAGT |
| | pcDNA3.1-GAP43-R | TCTTGTTGACTCCGAGCTCGGTACCAAGC |
| | TRPV1-F | CGAGCTCGGAGTCAACAAGATTGCACAAGAGAGCAAGA |
| | TRPV1-R | CTGCAGAATTTTTCTCCCCTGGGACCATGGA |
| Flag-TRPV1-GFP | pEGFP-N1-F | CCTTGTAATCTAGCGCTAGCGGATCTGACG |
| | pEGFP-N1-R | AGGGGAGAAAGAGCTCAAGCTTCGAATTCTGCAG |
| | TRPV1-F | GCTAGCGCTAGATTACAAGGATGACGACGATAAGATGGAACA |
| | TRPV1-R | GCTTGAGCTCTTTCTCCCCTGGGACCATGGAAT |
| GST-TRPV1 Ct Δ820 | pGEX-5X-1-F | TATACGGGAGAGCGGCCGCATCGT |
| | pGEX-5X-1-R | TCTTGTTGACTTCGGGGATCCCACGACCT |
| | TRPV1-F | GATCCCCGAAGTCAACAAGATTGCACAAGAGAGCAAG |
| | TRPV1-R | GCGGCCGCTCTCCCGTATAATGCTTCAGTTGAACTTCTTCC |
| GST-TRPV1 Ct | F | TTATACGGCCGCACTTAAGCCAGAGGATGCTGAGGTTTTCAAGG |
| | R | CTCTGGCTTAAGTGCCGCCGTATAATGCTTCAGTTGAACTTC |

| | | |
|---|---|---|
| GST-TRPV1 Ct (S818A, G819A, S820A) | F | AACTGAAGCATTATGCAGCCGCACTTAAGCCAGAGGATGCTG |
| | R | GGCTTAAGTGCGGCTGCATAATGCTTCAGTTGAACTTCTTCC |
| GST-TRPV1 Ct (Y817A, S818A, G819A, S820A) | F | AACTGAAGCATGCCGCAGCCGCACTTAAGCCAGAGGATGCTG |
| | R | GGCTTAAGTGCGGCTGCGGCATGCTTCAGTTGAACTTCTTCC |
| GST-TRPV1 Ct (S818A, G819A, S820A, L821A) | F | CAACTGAAGCATTATGCAGCCGCAGCCAAGCCAGAGGATGCTG |
| | R | CAGCATCCTCTGGCTTGGCTGCGGCTGCATAATGCTTCAGTTG |
